# Mapping the functional form of the trade-off between infection resistance and reproductive fitness under dysregulated immune signaling

**DOI:** 10.1101/2023.08.10.552815

**Authors:** Justin T. Critchlow, Arun Prakash, Katherine Y. Zhong, Ann T. Tate

## Abstract

Immune responses benefit organismal fitness by clearing parasites but also exact costs associated with immunopathology and energetic investment. Hosts manage these costs by tightly regulating the induction of immune signaling to curtail excessive responses and restore homeostasis. Despite the theoretical importance of turning off the immune response to mitigate these costs, experimentally connecting variation in the negative regulation of immune responses to organismal fitness remains a frontier in evolutionary immunology. In this study, we used a doseresponse approach to manipulate the RNAi-mediated knockdown efficiency of *cactus* (IκBα), a central regulator of Toll pathway signal transduction in flour beetles (*Tribolium castaneum*). By titrating *cactus* activity along a continuous gradient, we derived the shape of the relationship between immune response investment and traits associated with host fitness, including infection susceptibility, lifespan, fecundity, body mass, and gut homeostasis. *Cactus* knock-down increased the overall magintude of inducible immune responses and delayed their resolution in a dsRNA dose-dependent manner, promoting survival and resistance following bacterial infection. However, these benefits were counterbalanced by dsRNA dose-dependent costs to lifespan, fecundity, body mass, and gut integrity. Our results allowed us to move beyond the qualitative identification of a trade-off between immune investment and fitness to actually derive its functional form. This approach paves the way to quantitatively compare the evolution and impact of distinct regulatory elements on life-history trade-offs and fitness, filling a crucial gap in our conceptual and theoretical models of immune signaling network evolution and the maintenance of natural variation in immune systems.

## Introduction

An effective immune response is essential for recognizing and clearing infections. Hosts produce immune effectors even in the absence of infection as part of the constitutive response, but once they recognize invaders they inducibly create immune effectors to stymie the infection and then deactivate the response once the infection is under control during the decay or resolution phase of an immune response (Sheldon & Verhulst, 1996; Jent et al., 2019). Failure to manage infections can result in parasite-mediated pathology and exploitation, but excessive immune activation can also exert significant immunopathological and energetic costs (Graham et al., 2005; Sears et al., 2011; Badinloo et al., 2018). To balance these costs, hosts must tightly control immunological responses before, during, and after infections (Lazzaro & Tate, 2022).

By reducing the probability of colonization and initial replication by pathogens and other parasites, constitutive immune defenses mitigate the opportunity for parasite-induced pathology and discourage parasite transmission (Shudo & Iwasa, 2001; Westra et al., 2015; Boots & Best, 2018). Since constitutive immunity requires continuous resource investment, its benefit to fitness diminishes as the risk of infection declines (Hamilton et al., 2008). In contrast, inducible defenses are predicted to limit immunological costs in the absence of infection but risk being overwhelmed by fast-growing parasites if the response is not robust enough (Hamilton et al., 2008). In *Drosophila melanogaster* flies infected with the bacterium *Providencia rettgeri*, for example, host death is determined by whether the fly produces enough antimicrobial peptides (AMPs) before *P. rettgeri* replicates to uncontrollable numbers (Duneau et al., 2017). Yet, failure to dampen an induced immune response can also pose serious risks to the host’s health and lifespan, as seen when aged mealworm beetles overactivate the melanization response, damaging their Malpighian tubules (Khan et al., 2017). To prevent such runaway immunological costs, immune signaling pathways utilize negative feed-back loops (Ferrandon et al., 2007; Frank & Schmid-Hempel, 2019).

Robust negative regulation is predicted to affect the constitutive levels of immune proteins as well as the rate of induction, total magnitude, and decay rate of inducible immune responses (Figure 1) (Lazzaro & Tate, 2022). Most invertebrate species rely on regulatory proteins to tightly govern the recognition, signaling, and output of the NF-κB Imd and Toll pathways (Kleino & Silverman, 2014; F. Wang & Xia, 2018; Zhai et al., 2018; Valanne et al., 2022), and through genetic modification, researchers have started to characterize the consequences of their dysregulation (Lee & Ferrandon, 2011). For instance, the negative regulator Pirk interrupts the interaction of the transmembrane peptidoglycan recognition protein PGRP-LC and the intracellular signaling protein Imd (Aggarwal et al., 2008; Kleino et al., 2008; Basbous et al., 2011; Vincent & Dionne, 2021). When expression of *pirk* is disrupted via RNAi in the mosquito *Aedes aegypti*, survival against bacterial infection increases, but at a cost to female egg production (M. Wang et al., 2022). Cactus, the Toll inhibitor of nuclear factor kappa B (IκBα), prevents the translocation of the Toll pathway TFs Dif and Dorsal (Belvin & Anderson, 1996). Silencing and loss of function of the *cactus* gene in *D. melanogaster* disrupts Toll signaling resulting in increased production of AMPs and hemocyte proliferation (Qiu et al., 1998). While this Toll pathway dysregulation increases resistance and survival to infection (Lemaitre et al., 1996; Garver et al., 2009; Rhodes et al., 2018), it also shortens host lifespan (Ulrich et al., 2015), disrupts neuromuscular function (Beramendi et al., 2005), reduces lipid stores by suppressing insulin signaling (DiAngelo et al., 2009), and disrupts gut stability (Ryu et al., 2008). Unrestrained immune signaling clearly has consequences for host fitness, but how does every unit of gain in resistance translate into a cost to reproduction, and is the relationship linear or a case of diminishing returns?

**Figure 1.**
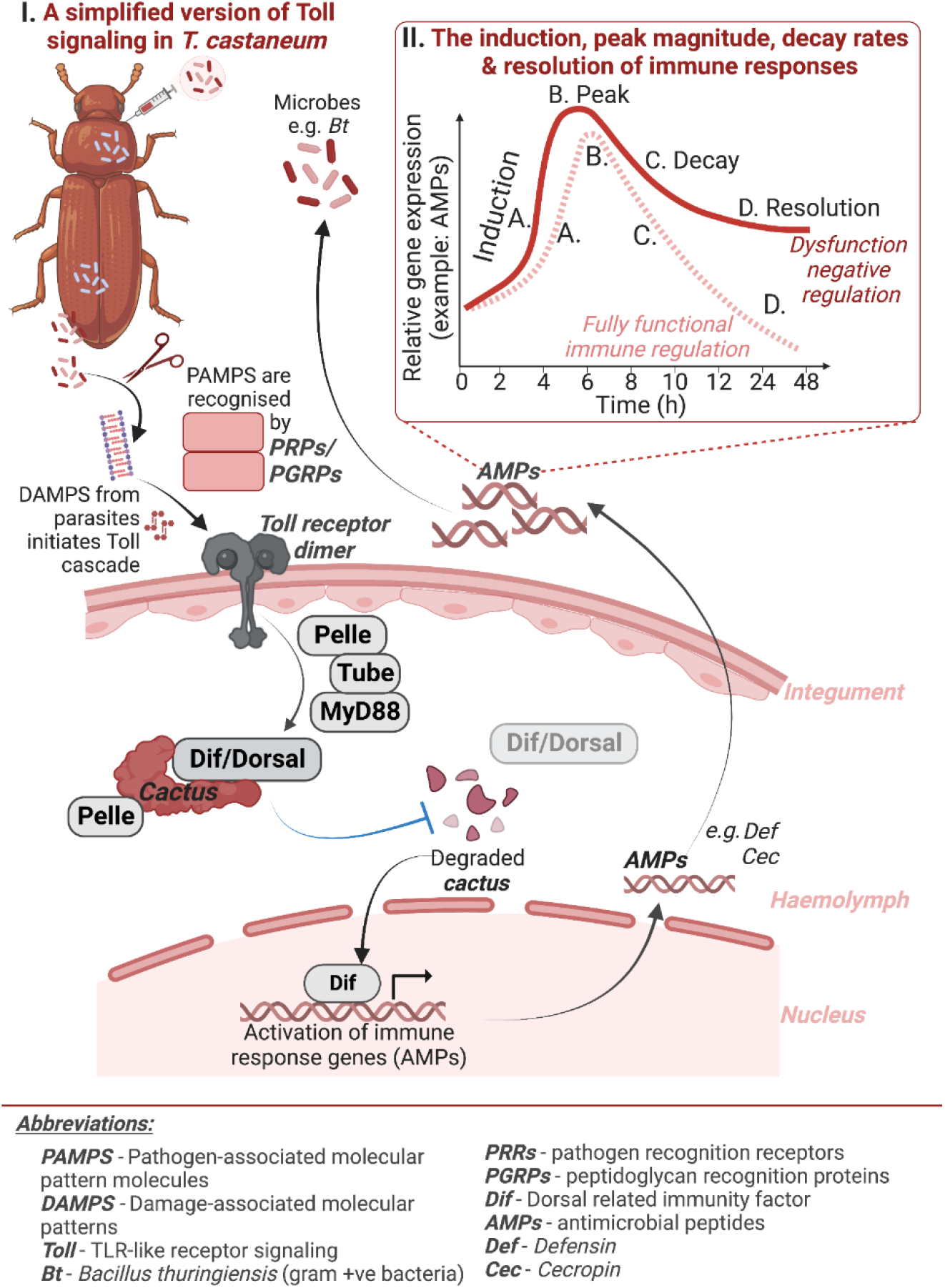
Simplified overview of the flour beetle Toll signaling pathway. **I)** Pattern Recognition Receptors (PRRs) identify pathogen or danger-associated molecular patterns (PAMPs or DAMPs) triggering signal transduction through Toll. This forms an intracellular signaling scaffold consisting of MyD88, Tube, and Pelle, resulting in Cactus phosphorylation by Pelle. The subsequent degradation of Cactus allows Dif and Dorsal transcription factors to translocate into the nucleus, initiating immune effector transcription. **II)** Production of AMPs from Toll signaling proceeds through a sequence of stages, starting with constitutive production before infection **(A)**, induction upon parasite recognition **(B)**, peak output **(C)**, decay of transcript production, and finally resolution of the response **(D)**. Dysfunction in negative regulators could potentially disturb signaling dynamics at any stage, leading to overproduction of immune effectors.

We do not yet have an answer to this question because previous studies have typically employed binary treatments where genes are either expressed normally or maximally disrupted through mutation, deletion, or efficient RNAi knock-down (Clayton et al., 2013; Hou et al., 2014; Prakash et al., 2021). While these approaches are critical for establishing the presence of a trade-off, dichotomous experimental designs limit our ability to quantitatively assess the impact of variation in immune dynamics on host-parasite coevolutionary dynamics and the costs associated with immune activation (Frank & Schmid-Hempel, 2019). By adopting an approach that titrates the magnitude of genetic manipulation, we can capture the continuous relationship between negative immune regulators and host fitness traits. This would enable us to mimic the natural variation commonly seen in immune gene expression and function and gain a deeper understanding of the evolutionary constraints and selection pressures imposed on regulatory immune genes.

In the present study, we developed an experimental framework to establish the functional form of the relationship between gains in infection resistance and costs to other life history traits for a specific regulatory node in immune signaling. We took advantage of dose-dependent RNAi knockdown efficiency in the red flour beetle (*Tribolium castaneum*) to quantitatively control the magnitude of dysregulation of Cactus. We measured the temporal dynamics of AMP transcription after exposure to fungal and bacterial microbes to parse apart the role of this individual regulator in fine tuning the rate of response to detection as well as the resolution of the response, both of which are predicted to determine an “optimal” immune response to infection (Frank, 2002). We also quantified functional metrics of infection susceptibility and resistance, including cellular immunity, hemolymph antimicrobial activity, bacterial load, and survival following live infection. At the same time, we quantified the impact of variation in Cactus activity on beetle fecundity, survival in the absence of infection, and gut homeostasis. Our results directly advance our understanding of immune system evolution and the complex feedbacks of host-parasite interactions on organismal fitness. These insights provide valuable information for understanding the pathogenesis of immune-related disorders and may guide the development of targeted therapeutic interventions, including siRNA-based therapies that aim to modulate immune responses for disease treatment.

## Results

### Cactus RNAi increases constitutive and total transcriptional activation and delays decay of Toll signaling

To investigate the role of *cactus* on *T. castaneum* immune signaling, we injected 250 ng of *cactus* or *malE* (control) dsRNA into adult beetles and then septically challenged them three days later, using either heat-killed *Bacillus thuringiensis* (Bt) or heat-killed *Candida albicans*, which contain different sets of microbe-associated molecular patterns (MAMPs) that potentially stimulate differences in recognition and subsequent immune signaling. We collected beetles at several time points over the next 48 hours, beginning before microbial challenge, and used RT-qPCR to measure *cactus* knockdown efficiency and the transcriptional induction and decay of three AMPs (Figure 2a-h) associated with Toll and IMD signaling (*cecropin-2* (Toll), *defensin-2* (IMD), and *defensin-3* (Toll & IMD) (Yokoi, Koyama, Minakuchi, et al., 2012; Herndon et al., 2020)). We expected that our control (MalE) beetles would follow typical temporal dynamics (Figure 1), where AMP transcript abundance reaches maximum levels in 4-12 hours post septic challenge, then returns to baseline by 48 hours (Zou et al., 2007; Tate & Graham, 2017).

**Figure 2.**
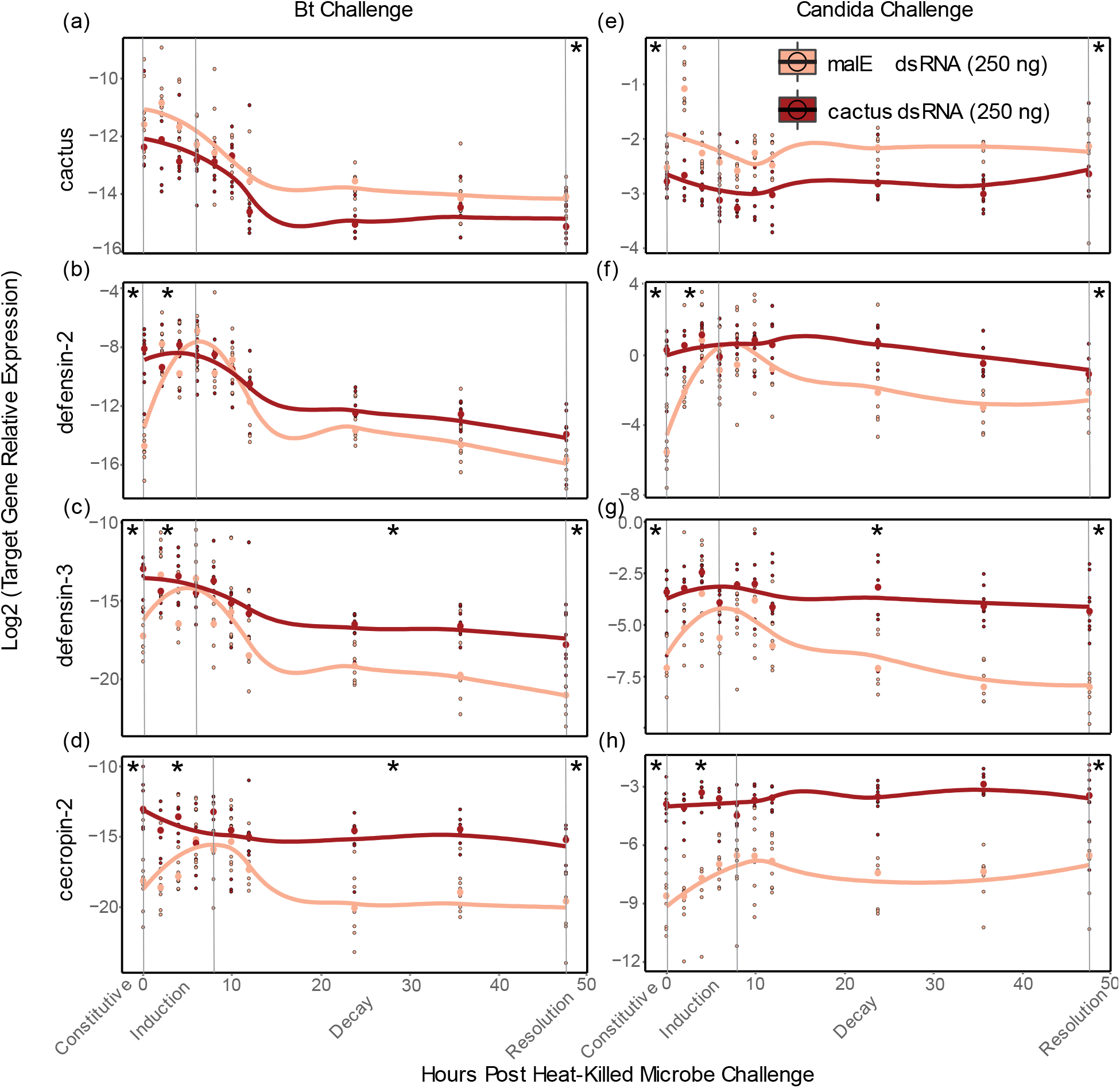
DsRNA-mediated knockdown of *cactus* results in increased Toll signaling. **a-h)** The expression of *cactus* **(a, e)** and the antimicrobial peptides *defensin-2* **(b, f)**, *defensin-3* **(c, g)** and *cecropin-2* **(d, h)** were assayed via RT-qPCR in whole adult beetles treated with 250 ng of *cactus* or *malE* dsRNA and then septically challenged with heat-killed Bt (left) or *C. albicans* (right). Beetles were sacrificed before microbial challenge (hour 0), and nine additional times after challenge over 48 hours. The expression of each gene relative to the reference gene RP18s is represented on a log2 scale. Splines have been added to visualize the induction and decay dynamics from RNAi treatment using “loess” function (span 0.25) in the geom_smooth algorithm of ggplot2 in R. The * represents whether the constitutive (hour 0), induction (slope from hour 0 to peak expression), decay (slope from peak expression to 48 hours), or resolution (hour 48) windows were significantly altered by *cactus* depletion (α = 0.0167).

Therefore, to statistically analyze differences in temporal dynamics among treatments, we considered four outputs for AMP expression: constitutive (hour 0), the rate of induction, the rate of decay from the peak magnitude of expression, and the magnitude of AMP production when the response should be resolved (hour 48).

*Cactus* RNAi treatment significantly reduced total *cactus* expression, relative to MalE RNAi-treated beetles, across the time course of the Bt (*p* < .001, Table 1) and *Candida* exposure experiments (*p* < 2 x 10^-16^, Table 2). Averaged across all time points, relative transcript abundance decreased by 39% (Bt) and 33% (*Candida*). For all three AMPs, constitutive expression and the overall magnitude of transcription after exposure were higher in *cactus-* depleted beetles, independent of the challenged microbe (Figure 2b-d & 2f-h) (Tables 1 & 2). AMP induction rates were significantly higher for *def-2*, *def-3*, and *cec-2* when challenged with Bt (*p <* 0.01) and for *def-2* and *cec-2* when challenged with *Candida* (*p* < 0.01). Additionally, *cactus* RNAi delayed decay for *def-3* and *cec-2* when challenged with Bt (*p* < 0.01), and delayed decay for *def-3* when challenged with *Candida* (*p <* 0.01). Resolution transcription levels at 48 hours post exposure were significantly higher for *def-3* and *cec-2* (*p* < 0.001) but not *def-2* (*p =* 0. 03) when challenged with Bt and were higher for *def-3, cec-2*, and *def-2* (*p* < 0.01) when challenged with *Candida*.

**Table 1.**
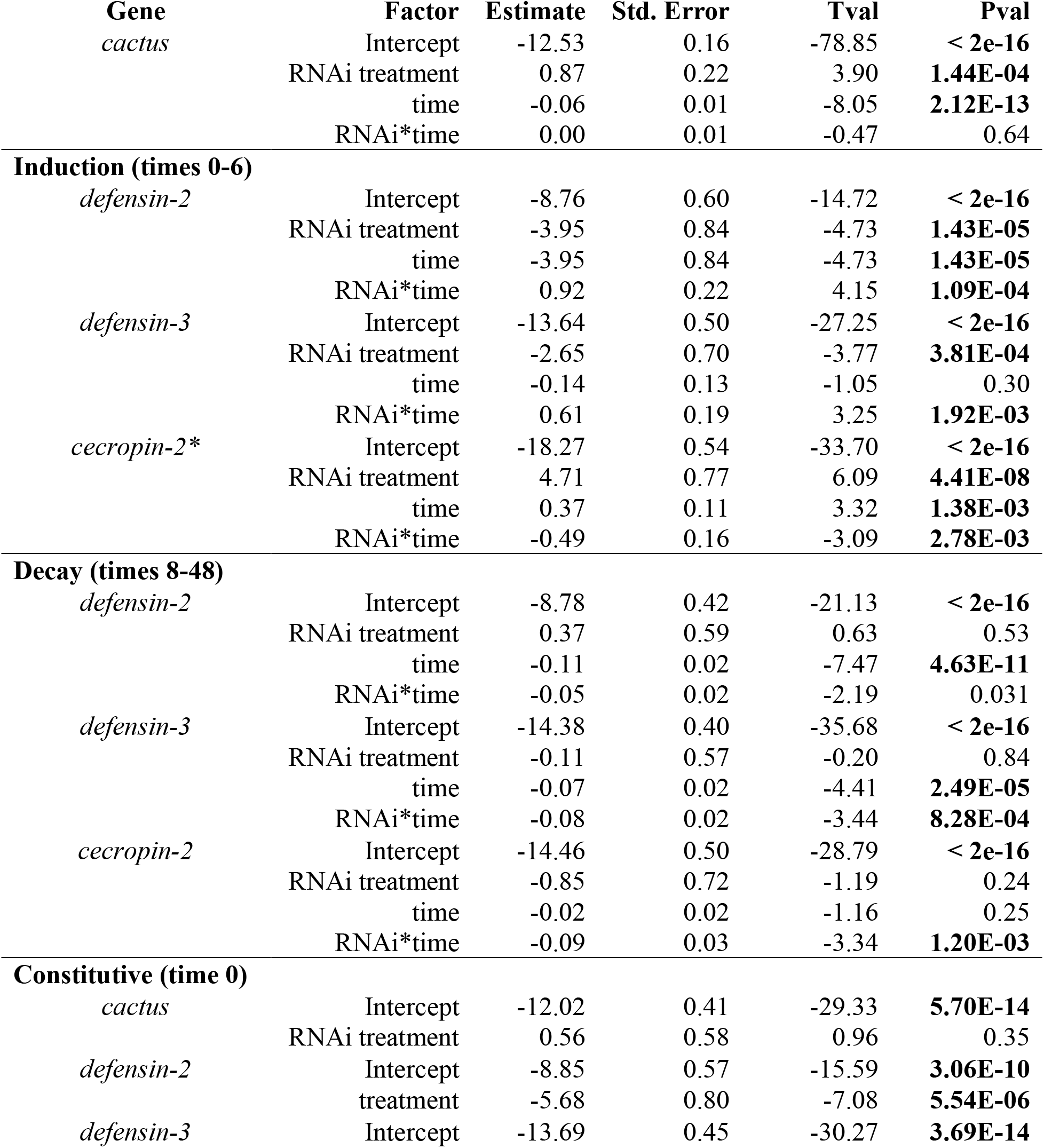
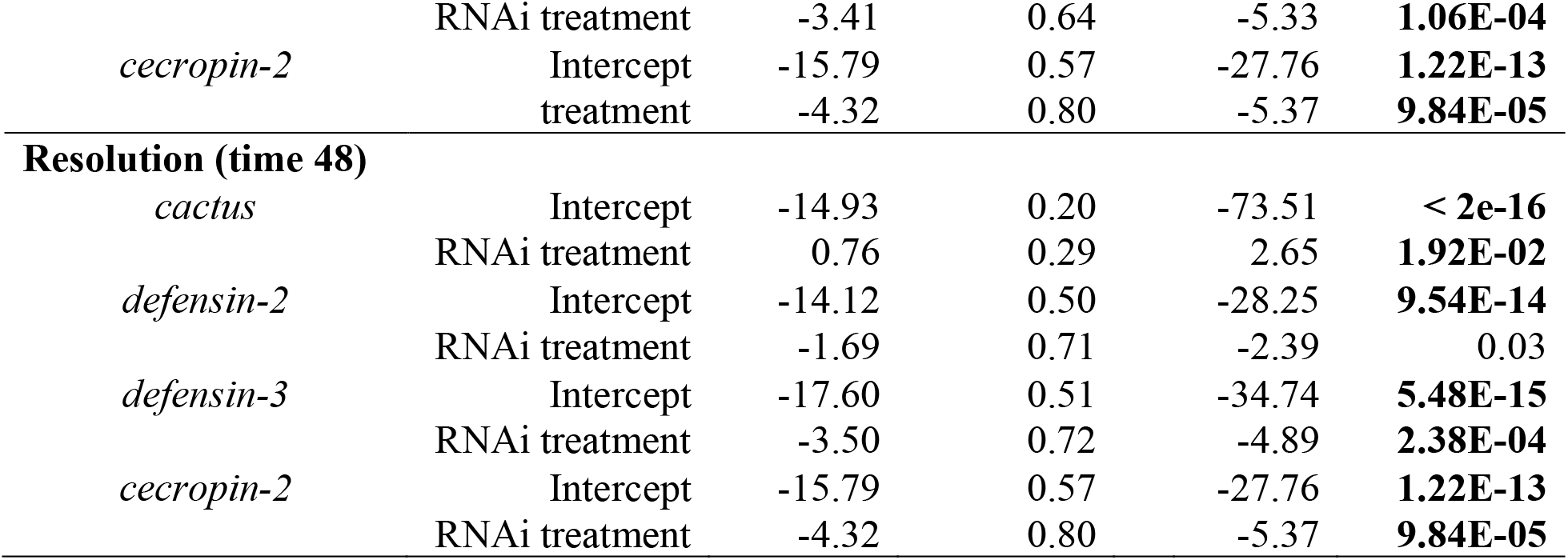
Effect of *cactus* RNAi (250ng) on knockdown efficiency and AMP gene expression to *Bacillus thuringiensis* infection in flour beetles compared to MalE control. Linear models (expression ∼ RNAi treatment*time) conducted with “lm” function in R. Time denotes the hours passed post microbial challenge. Expression values are on a log scale. For AMP genes, P values less than the significance threshold after correcting for multiple testing (Bonferroni method; α = 0.0167) are in bold. * Induction for cecropin-2 ends at 8 times post infection.

**Table 2.**
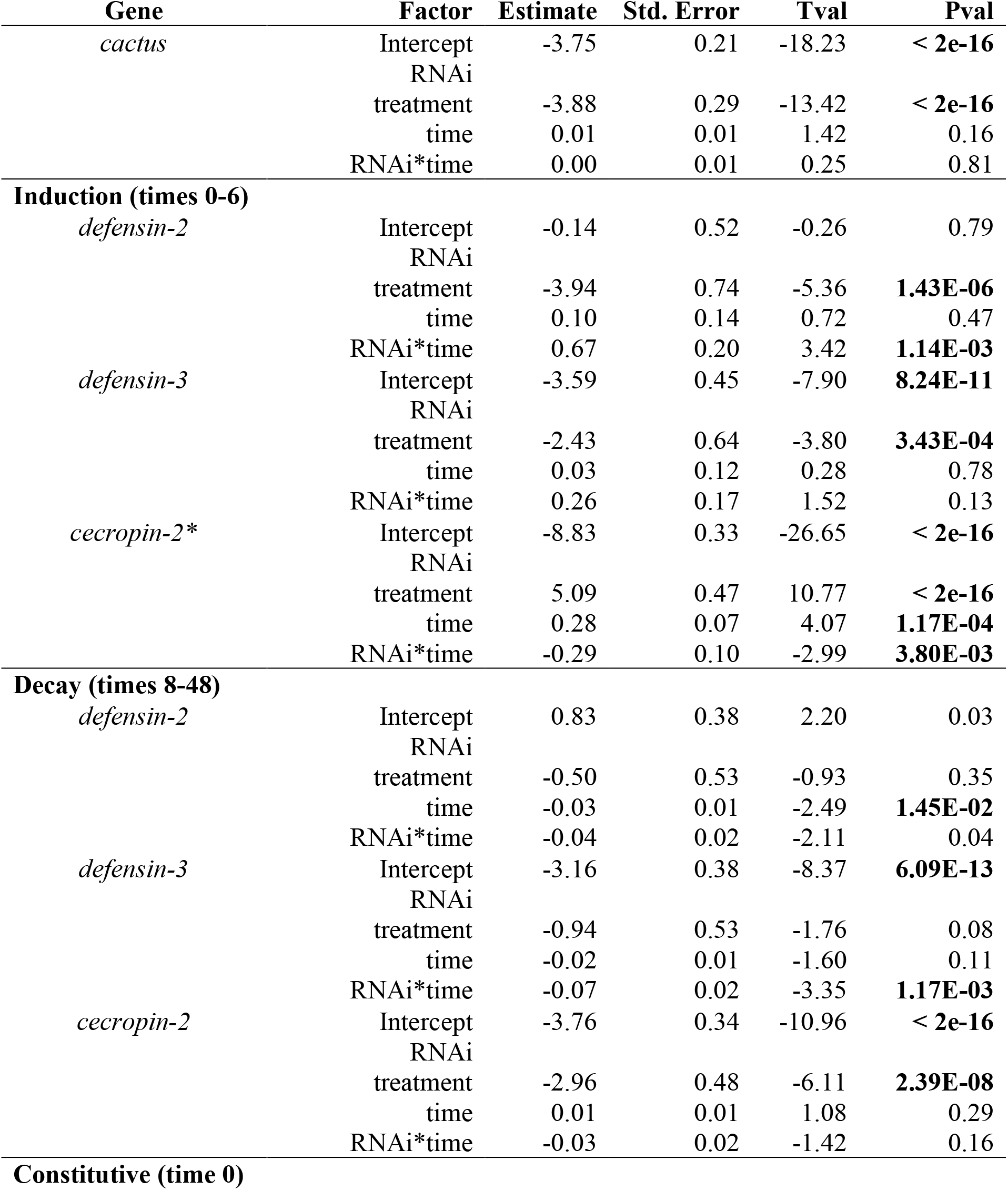
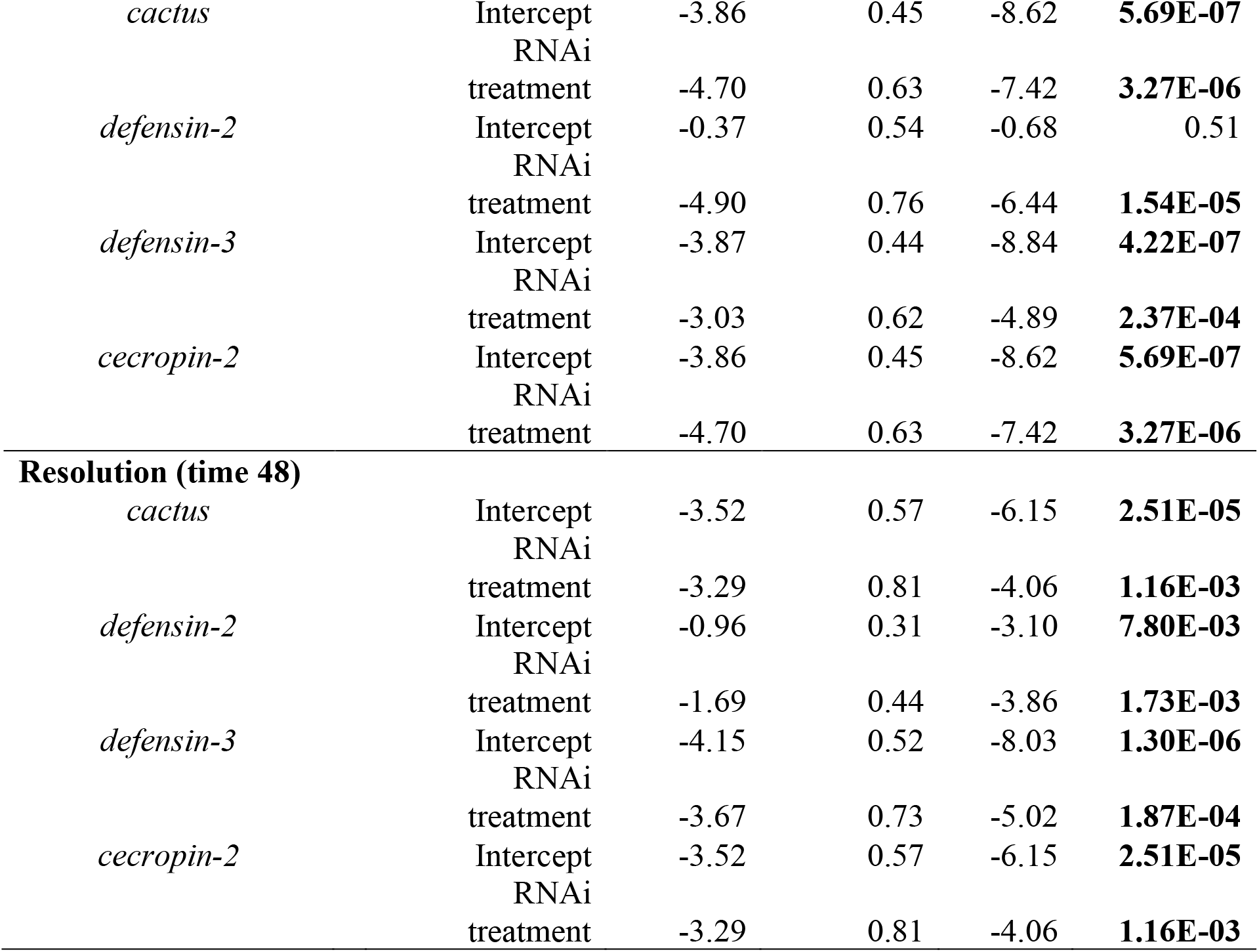
Effect of *cactus* RNAi (250ng) on knockdown efficiency and AMP gene expression to *Candida albicans* infection in flour beetles compared to MalE control. Linear models (expression ∼ RNAi treatment*time) conducted with “lm” function in R. Time denotes the hours passed post microbial challenge. Expression values are on a log scale. For AMP genes, P values less than the significance threshold after correcting for multiple testing (Bonferroni method; α = 0.0167) are in bold. * Induction for cecropin-2 ends at 8 times post infection.

| Gene | Factor | Estimate | Std. Error | Tval | Pval |
| --- | --- | --- | --- | --- | --- |
| *cactus* | Intercept | -3.75 | 0.21 | -18.23 | **< 2e-16** |
|  | RNAi |  |  |  |  |
|  | treatment | -3.88 | 0.29 | -13.42 | **< 2e-16** |
|  | time | 0.01 | 0.01 | 1.42 | 0.16 |
|  | RNAi*time | 0.00 | 0.01 | 0.25 | 0.81 |
| **Induction (times 0-6)** |  |  |  |  |  |
| *defensin-2* | Intercept | -0.14 | 0.52 | -0.26 | 0.79 |
|  | RNAi |  |  |  |  |
|  | treatment | -3.94 | 0.74 | -5.36 | **1.43E-06** |
|  | time | 0.10 | 0.14 | 0.72 | 0.47 |
|  | RNAi*time | 0.67 | 0.20 | 3.42 | **1.14E-03** |
| *defensin-3* | Intercept | -3.59 | 0.45 | -7.90 | **8.24E-11** |
|  | RNAi |  |  |  |  |
|  | treatment | -2.43 | 0.64 | -3.80 | **3.43E-04** |
|  | time | 0.03 | 0.12 | 0.28 | 0.78 |
|  | RNAi*time | 0.26 | 0.17 | 1.52 | 0.13 |
| *cecropin-2\** | Intercept | -8.83 | 0.33 | -26.65 | **< 2e-16** |
|  | RNAi |  |  |  |  |
|  | treatment | 5.09 | 0.47 | 10.77 | **< 2e-16** |
|  | time | 0.28 | 0.07 | 4.07 | **1.17E-04** |
|  | RNAi*time | -0.29 | 0.10 | -2.99 | **3.80E-03** |
| **Decay (times 8-48)** |  |  |  |  |  |
| *defensin-2* | Intercept | 0.83 | 0.38 | 2.20 | 0.03 |
|  | RNAi |  |  |  |  |
|  | treatment | -0.50 | 0.53 | -0.93 | 0.35 |
|  | time | -0.03 | 0.01 | -2.49 | **1.45E-02** |
|  | RNAi*time | -0.04 | 0.02 | -2.11 | 0.04 |
| *defensin-3* | Intercept | -3.16 | 0.38 | -8.37 | **6.09E-13** |
|  | RNAi |  |  |  |  |
|  | treatment | -0.94 | 0.53 | -1.76 | 0.08 |
|  | time | -0.02 | 0.01 | -1.60 | 0.11 |
|  | RNAi*time | -0.07 | 0.02 | -3.35 | **1.17E-03** |
| *cecropin-2* | Intercept | -3.76 | 0.34 | -10.96 | **< 2e-16** |
|  | RNAi |  |  |  |  |
|  | treatment | -2.96 | 0.48 | -6.11 | **2.39E-08** |
|  | time | 0.01 | 0.01 | 1.08 | 0.29 |
|  | RNAi*time | -0.03 | 0.02 | -1.42 | 0.16 |
| **Constitutive (time 0)** |  |  |  |  |  |
| *cactus* | Intercept | -3.86 | 0.45 | -8.62 | **5.69E-07** |
|  | RNAi |  |  |  |  |
|  | treatment | -4.70 | 0.63 | -7.42 | **3.27E-06** |
| *defensin-2* | Intercept | -0.37 | 0.54 | -0.68 | 0.51 |
|  | RNAi |  |  |  |  |
|  | treatment | -4.90 | 0.76 | -6.44 | **1.54E-05** |
| *defensin-3* | Intercept | -3.87 | 0.44 | -8.84 | **4.22E-07** |
|  | RNAi |  |  |  |  |
|  | treatment | -3.03 | 0.62 | -4.89 | **2.37E-04** |
| *cecropin-2* | Intercept | -3.86 | 0.45 | -8.62 | **5.69E-07** |
|  | RNAi |  |  |  |  |
|  | treatment | -4.70 | 0.63 | -7.42 | **3.27E-06** |
| **Resolution (time 48)** |  |  |  |  |  |
| *cactus* | Intercept | -3.52 | 0.57 | -6.15 | **2.51E-05** |
|  | RNAi |  |  |  |  |
|  | treatment | -3.29 | 0.81 | -4.06 | **1.16E-03** |
| *defensin-2* | Intercept | -0.96 | 0.31 | -3.10 | **7.80E-03** |
|  | RNAi |  |  |  |  |
|  | treatment | -1.69 | 0.44 | -3.86 | **1.73E-03** |
| *defensin-3* | Intercept | -4.15 | 0.52 | -8.03 | **1.30E-06** |
|  | RNAi |  |  |  |  |
|  | treatment | -3.67 | 0.73 | -5.02 | **1.87E-04** |
| *cecropin-2* | Intercept | -3.52 | 0.57 | -6.15 | **2.51E-05** |
|  | RNAi |  |  |  |  |
|  | treatment | -3.29 | 0.81 | -4.06 | **1.16E-03** |

### Amplification of Toll Signaling via cactus RNAi increases total circulating hemocytes

To study the impact of enhanced Toll signaling on cellular immunity, we measured circulating hemocyte counts in adult beetles before and after heat-killed Bt challenge (Figure 3a). Circulating hemocytes did not constitutively increase in c*actus* RNAi (250 ng) treated beetles before Bt challenge or after a sham infection (Table S1). However, *cactus* RNAi-treated beetles exhibited significantly higher numbers of hemocytes at all measured time points after bacterial exposure relative to their time-matched MalE counterparts (Wilcoxon rank sum test and FDR correction; Table S1).

**Figure 3.**
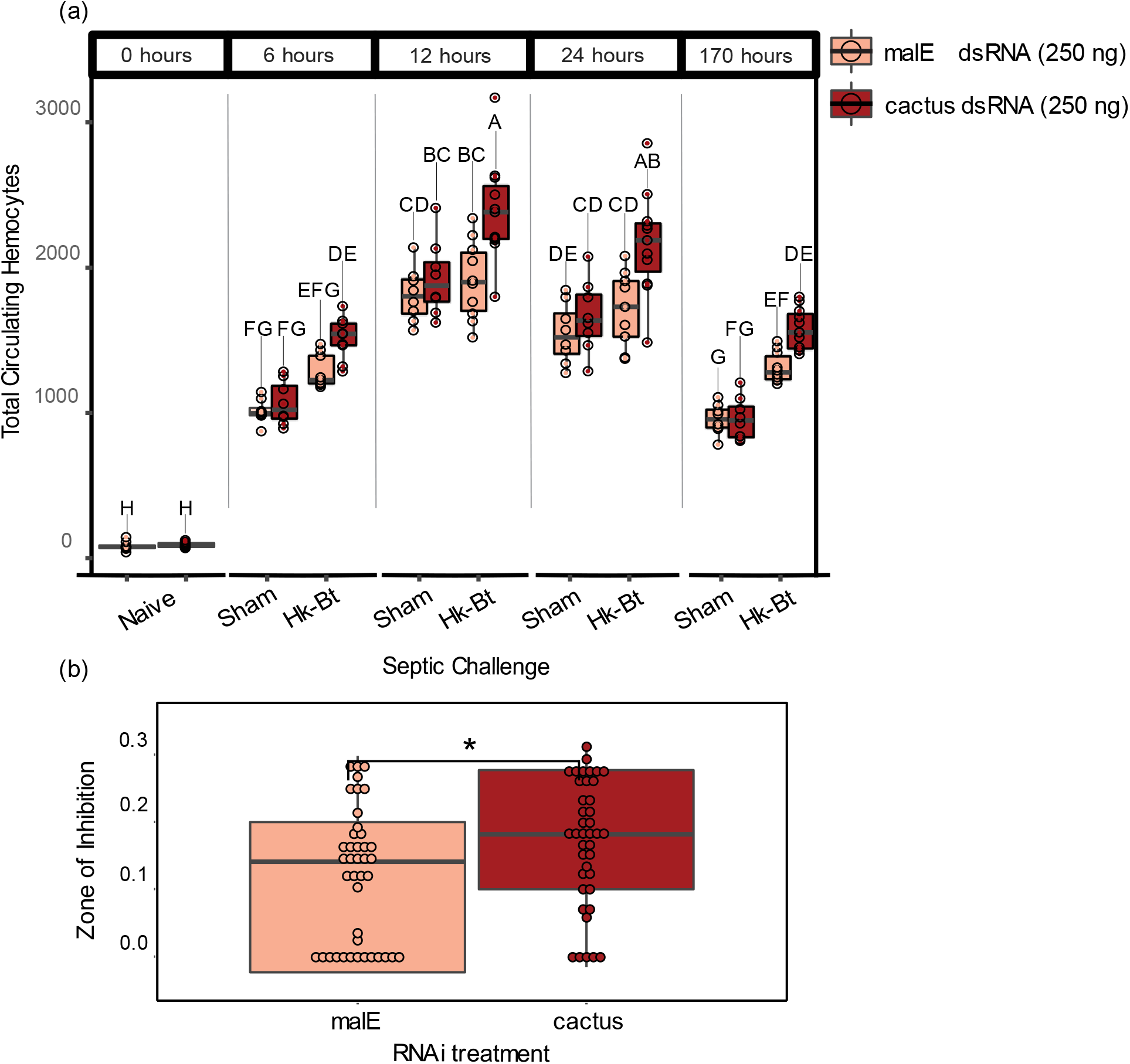
Toll signaling increases functional metrics of cellular immunity and antibacterial activity **a)** The number of circulating hemocytes in malE or cactus dsRNA-treated adult beetles three days after dsRNA exposure, before (naïve) or after subsequent exposure to a sterile saline (sham) or heat-killed *Bacillus thuringiensis* (hk-Bt). Groups not sharing the same letter are significantly different (Wilcoxon rank sum test and FDR correction, P < 0.05). **b)** The antibacterial activity of whole-beetle homogenates was assessed by measuring the mean diameter in mm of the zone of inhibition of bacterial growth on agar plates (* indicates p < 0.05).

### Uninhibited Toll signaling enhances antibacterial activity

To measure whether unrestrained Toll signaling correlates with functional changes in antibacterial activity, we used a lytic zone assay to quantify the ability of beetle hemolymph to prevent Bt growth on Luria Broth agar (Figure 3b). Beetles treated with *cactus* dsRNA exhibited significantly increased hemolymph antibacterial activity relative to MalE controls (Kruskal-Wallace rank sum test, N = 40-41 beetles/treatment, (χ2 = 6.08, Df = 1, *P =* 0.014).

### The magnitude of Toll pathway activation depends on the dose of cactus dsRNA

To quantify the relationship between the magnitude of immune dysregulation and immune dynamics, we repeated the previous experiment but provided beetles with one of four *cactus* dsRNA doses: 0 ng (but 250 ng of *malE* dsRNA), 2.5 ng, 25 ng, or 250 ng. Using the linear model (expression ∼ dose), the dose of *cactus* dsRNA was significantly associated with decreased total *cactus* transcription (*p* < 0.001) (Figure 4a, Table 3). Total relative *cactus* transcript abundance was reduced by 25% (2.5 ng), 32% (25 ng), and 35% (250 ng) for each respective dose.

**Figure 4.**
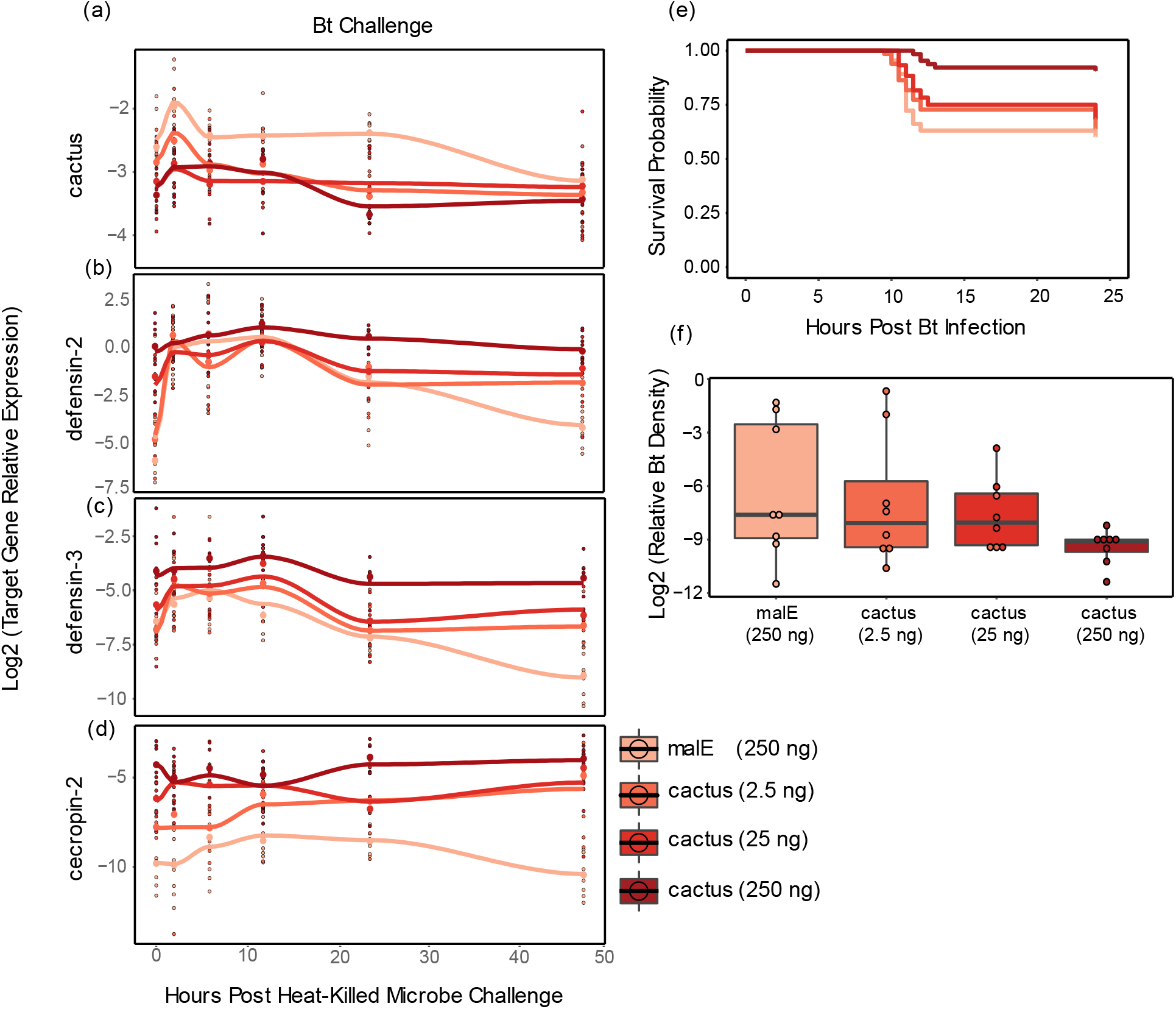
Quantitative knockdown of cactus transcripts benefits resistance and survival during infection. **a-d)** The expression of *cactus* **(a)** and the antimicrobial peptides *defensin-2* **(b)**, *defensin-3* **(c)** and *cecropin-2* **(d)** were assayed via qRT-PCR in whole adult beetles treated with 250, 25, or 2.5 ng of *cactus dsRNA* or 250 ng of *malE* dsRNA and then septically challenged with heat-killed Bt. Beetles were frozen before microbial challenge, time 0, and five additional times over 48 hours. Splines have been added to visualize the induction and decay dynamics from RNAi treatment using “loess” function in the geom_smooth algorithm of ggplot2 (R). See Table 3 for statistical analyses, which are not visualized here due to complexity. **e)** Survival to an LD 50 (6.5 x 10^8/mL) Bt infection after RNAi treatment was monitored for 24 hours (N = 60-64 beetles/treatment and 29-34 per sex). **f)** To measure shifts in host resistance to bacterial infection from *cactus* RNAi treatment, beetles were given an LD-50 dose of Bt and sacrificed seven hours later. Relative bacterial density for each individual within each dsRNA treatment was quantified via RT-qPCR and calculated as the difference between Bt-specific and host reference gene expression (RP18s) on a log2 scale.

**Table 3.** Effect of *cactus* RNAi dose on knockdown efficiency and AMP gene expression to *Bacillus thuringiensis* infection in flour beetles compared to MalE control. Linear models (expression ∼ dose*time) conducted with “lm” function in R. Time denotes the hours passed post microbial challenge while dose represents the effect of *cactus* dsRNA per ng. Expression values are on a log scale. P values less than the significance threshold after correcting for multiple testing (Bonferroni method; α = 0.0167) are in bold.

| Gene | Factor | Estimate | Std. Error | t value | Pval |
| --- | --- | --- | --- | --- | --- |
| *cactus* | Intercept | -2.64 | 0.06 | -46.06 | **< 2e-16** |
|  | dose | 0.00 | 0.00 | -3.61 | **3.89E-04** |
|  | time | -0.01 | 0.00 | -4.13 | **5.50E-05** |
|  | dose*time | 0.00 | 0.00 | 0.27 | 0.79 |
| **Induction (times 0-6)** |  |  |  |  |  |
| *defensin-2* | Intercept | -2.70 | 0.36 | -7.44 | **5.20E-11** |
|  | dose | 0.01 | 0.00 | 3.60 | **5.23E-04** |
|  | time | 0.48 | 0.10 | 4.78 | **6.51E-06** |
|  | dose*time | 0.00 | 0.00 | -1.85 | 0.07 |
| *defensin-3* | Intercept | -6.24 | 0.24 | -25.82 | **< 2e-16** |
|  | dose | 0.01 | 0.00 | 3.90 | **1.85E-04** |
|  | time | 0.22 | 0.07 | 3.30 | **1.38E-03** |
|  | dose*time | 0.00 | 0.00 | -1.27 | 0.21 |
| *cecropin-2* | Intercept | -8.13 | 0.34 | -23.69 | **< 2e-16** |
|  | dose | 0.01 | 0.00 | 5.26 | **9.37E-07** |
|  | time | 0.10 | 0.09 | 1.04 | 0.30 |
|  | dose*time | 0.00 | 0.00 | -0.92 | 0.36 |
| **Decay (times 12-48)** |  |  |  |  |  |
| *defensin-2* | Intercept | 0.73 | 0.35 | 2.12 | 0.04 |
|  | dose | 0.00 | 0.00 | 0.66 | 0.51 |
|  | time | -0.07 | 0.01 | -6.71 | **1.71E-09** |
|  | dose*time | 0.00 | 0.00 | 2.16 | 0.03 |
| *defensin-3* | Intercept | -4.96 | 0.37 | -13.55 | **<2e-16** |
|  | dose | 0.01 | 0.00 | 1.84 | 0.07 |
|  | time | -0.06 | 0.01 | -4.86 | **4.93E-06** |
|  | dose*time | 0.00 | 0.00 | 1.33 | 0.19 |
| *cecropin-2* | Intercept | -6.89 | 0.51 | -13.53 | **<2e-16** |
|  | dose | 0.01 | 0.00 | 1.27 | 0.21 |
|  | time | -0.01 | 0.02 | -0.63 | 0.53 |
|  | dose*time | 0.00 | 0.00 | 1.48 | 0.14 |
| **Constitutive (time 0)** |  |  |  |  |  |
| *cactus* | Intercept | -2.82 | 0.10 | -28.85 | **<2e-16** |
|  | dose | 0.00 | 0.00 | -2.22 | **3.41E-02** |
| *defensin-2* | Intercept | -3.87 | 0.43 | -9.02 | **4.76E-10** |
|  | dose | 0.01 | 0.00 | 4.39 | **1.30E-04** |
| *defensin-3* | Intercept | -6.67 | 0.29 | -22.98 | **<2e-16** |
|  | dose | 0.01 | 0.00 | 3.84 | **5.99E-04** |
| *cecropin-2* | Intercept | -8.19 | 0.34 | -24.41 | **<2e-16** |
|  | dose | 0.02 | 0.00 | 5.94 | **1.67E-06** |
| **Resolution (time 48)** |  |  |  |  |  |
| *cactus* | Intercept | -3.22 | 0.09 | -34.65 | **<2e-16** |
|  | dose | 0.00 | 0.00 | -1.14 | 0.26 |
| *defensin-2* | Intercept | -2.52 | 0.31 | -8.23 | **3.47E-09** |
|  | dose | 0.01 | 0.00 | 4.08 | **3.10E-04** |
| *defensin-3* | Intercept | -7.41 | 0.33 | -22.64 | **<2e-16** |
|  | dose | 0.01 | 0.00 | 4.19 | **2.25E-04** |
| *cecropin-2* | Intercept | -7.33 | 0.53 | -13.83 | **1.49E-14** |
|  | dose | 0.01 | 0.00 | 3.16 | **3.60E-03** |

Expression of the three AMPs significantly increased with each increasing dose during the constitutive, induction, and resolution windows (Figure 4b-d, Table 3). For example, for every ng of *cactus* dsRNA, constitutive expression of *def-2* increased by 0.01 (*p* < 0.001), *def-3* increased by 0.01 (*p* < 0.001), and *cec-2* increased by 0.02 (*p* < 0.0001). Interestingly, non-significant interaction effects between dose and time indicate that increasing *cactus* dsRNA dose did not significantly alter induction and decay rates for the AMPs (Table 3). This result is most likely due to the 2.5 ng, 25 ng, and 250 ng treatments converging upon maximum expression during the acute infection phase, since increasing *cactus* dsRNA dose significantly increases AMP expression during the constitutive and resolution hours for all three AMPs.

### Elevated Toll pathway signaling increases survival to septic bacterial infection

We conducted a survival analysis to evaluate the relationship between *cactus* dsRNA dose and survival after live Bt infection. We administered one of four *cactus* dsRNA doses (N=60-65 per dose). After three days, beetles were septically challenged with an LD50 dose of Bt, which typically kills beetles within 12 hours. Weighted Cox Regression analysis revealed that for every ng of *cactus* dsRNA injected, the mortality risk from Bt declines by 0.69% (Figure 4e, Coef = - 0.01, Se(coef) = 2 x 10^-3^, hazard ratio = 0.993, lower 95% CI = 0.990, upper 95% CI = 0.997, Z = -3.94, *p* < 8.1 x 10^-5^). Furthermore, we assessed the impact of *cactus* dsRNA dose on *in vivo* bacterial loads at seven hours post-infection (Figure 4f). Since the data was non-normally distributed, we used the Kruskal-Wallace rank sum test, which did not reveal a significant difference between the *cactus* doses (χ2 = 4.30, Df = 3, *P =* 0.23). However, like previous experiments, our data displayed a bifurcating distribution of low and high Bt loads, complicating the statistical analysis (Duneau et al., 2017; Jent et al., 2019). When we repeated the experiment for 0 and 250 ng treatments to assay bacterial loads at a slightly earlier time point (6 hours), we observed a significant reduction in Bt load in the 250 ng treated beetles (estimate = -2.02, st. error = 0.84, t = -2.41, *p =* 0.03) (Figure S1).

### Increased Toll pathway signaling has severe fitness related costs

To assess the potential costs of enhanced Toll signaling, we measured changes to beetle lifespan, female egg laying, body mass, and gut integrity in response to all four *cactus* RNAi doses. We determined lifespan by monitoring beetle survival following RNAi injection, using the experimental setup from Figure 4e but without Bt infection (N = 30-32 beetles/dose, N=15-16 beetles/sex, Figure 5a). We then conducted a Weighted Cox Regression analysis and found that for every ng of *cactus* dsRNA the background mortality risk significantly increased by 0.89% (Table S2, hazard ratio = 1.01, Z = 16.25, *p* < 10^-8^). Additionally, our analysis revealed that male beetles had a significantly lower survival rate compared to female beetles (hazard ratio = 1.30, Z = 16.25, *p* < 10^-8^). We did not investigate this further as sex did not prove to be a significant factor in any other experiments.

**Figure 5.**
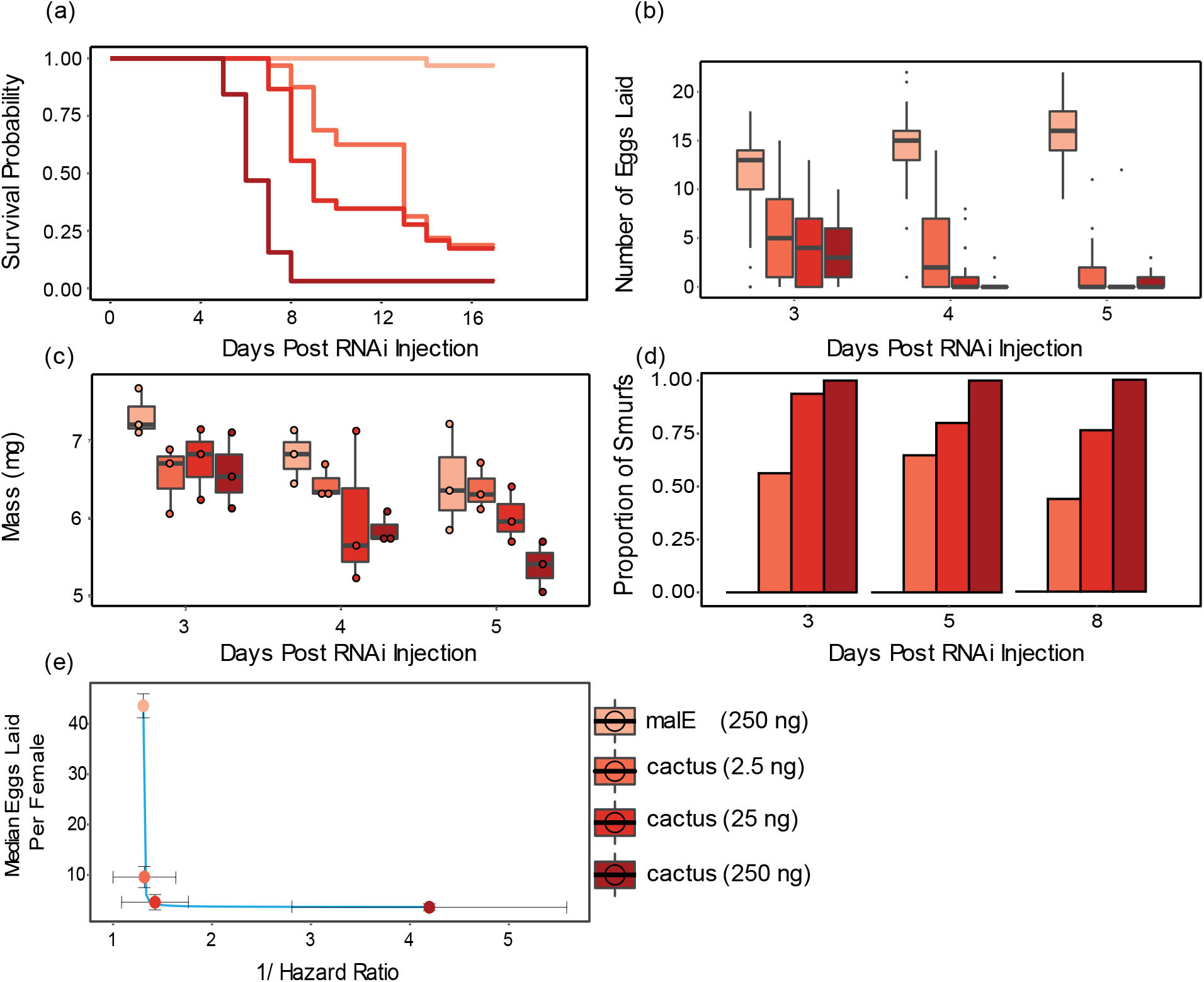
The costs of immune over-activation to fitness-associated traits. **a)** Beetle life-span after RNAi treatment was measured by monitoring survival in beetles given 250, 25, or 2.5 ng of *cactus dsRNA* or 250 ng of *malE* dsRNA for 17 days (N = 30-32 beetles/treatment and 15-16 beetles/sex). **b)** Female reproductive output after RNAi treatment was measured by allowing RNAi treated virgin female beetles 24 hours to mate and counting their number of eggs laid for three days. **c)** Female beetle mass after RNAi treatment was measured three to five days after *cactus* RNAi by pooling 3 individual beetles per measurement. Weighed beetles were discarded after measurement each day. **d)** Beetle gut integrity was measured by feeding adults flour stained blue and observing whether the blue dye entered the beetle hemolymph (N=15-21 beetles/day and 7-11 beetles/sex per day. **e)** The relationship between infection survival and fecundity (y = 1 / (-8.1 + 8.1 * x) + 3.0). Survival rate to infection for each RNAi treatment was calculated as 1/ (the hazard ratio relative to MalE). Reproductive output for each RNAi treatment is the median number of eggs laid for all three days measured per female.

Next, we investigated the impact of increased Toll signaling on female beetle reproductive output by measuring the number of eggs laid over consecutive 24 hour periods by females previously injected with *cactus* or *malE* dsRNA and then mated. Because the data is heteroscedastic, non-normal, and has an excess of 0 values, we analyzed the data using a Zero-Inflated Negative Binomial (ZINB) generalized linear model (dose + day + (1 | id)), which revealed that dose significantly reduced female egg laying (estimate = -3.57 x 10^-3^, st. error = 7.4 x 10^-4^, t = -4.86, *p* < 1.1 x 10^-6^) (Figure 5b, Table S3).

Since Toll pathway activation is known to disrupt insulin signaling resulting in reduced larval and adult weights (DiAngelo et al., 2009; Suzawa et al., 2019), we investigated the metabolic demands of increasing Toll signaling on female beetle body mass. Beetles were given one of four *cactus* RNAi doses and weighed on day three, four, or five. Using the linear model (mass ∼ dose + day), we found that *cactus* RNAi significantly decreased female body mass per ng of *cactus* dsRNA (estimate = -2.56 x 10^-3^, st. error = -7.94 x 10^-4^, t = -3.23, *p* < 0.01) (Figure 5c, Table S4). Furthermore, our analysis indicated that beetles weighed significantly less at later dates regardless of treatment (estimate = -0.38, std error = 0.10, t = -3.63, *p* < 0.001). These results coincide with our qualitative observations that the fat bodies of *cactus* treated beetles are depleted compared to MalE controls by day five.

In *D. melanogaster*, overactivation of AMPs exacerbates gut dysfunction, correlating with reduced lifespan (Rera et al., 2012). We investigated if Toll signaling dysfunction in beetles affects gut stability and if the severity of this dysfunction relates to gut instability. Beetles were given one of four *cactus* RNAi treatments and fed a non-absorbable blue food dye. If gut integrity is disrupted, the blue dye leaks into the hemolymph, causing the beetles to turn blue, which are called “smurfs” (Figure S2) (Rera et al., 2011). Analysis using a binomial glm (smurfs ∼ dose*day) showed that the dose of *cactus* dsRNA significantly influenced the proportion of smurf beetles (estimate = 0.18, st. error = 0.06, t = 3.13, *p* < 0.01) (Figure 5d). However, the number of days post RNAi injection and the interaction of dose and either day did not significantly influence the proportion of smurf beetles (Table S5).

## Discussion

In this study, we implemented a novel experimental framework to map the fitness-related benefits and costs of failing to properly regulate an immune signaling pathway. Poorly restrained Toll pathway activation in *T. castaneum* resulted in increased constitutive and inducible expression of AMPs, delayed transcriptional decay of AMPs during the resolution of the response, and increased numbers of microbe-induced circulating hemocytes. By serially modulating the dose of *cactus* dsRNA, we manipulated the magnitude of Toll signaling, allowing us to extract the functional form of the relationship between increased parasite resistance via humoral and cellular immunity and fitness-related traits. This analysis revealed a steep convex relationship between the benefits of immune investment against infection and the costs to fitness-associated traits like reproduction, reflecting the importance of Cactus as a central node in immune signaling as well as the pleiotropic impact of Toll pathway signaling on physiological homeostasis in insects. This experimental design can serve as a valuable blueprint for future studies to quantify the relationship between immune investment and reproductive fitness or to compare the relative roles of different regulatory checkpoints for the cost-benefit calculus of immune response regulation.

### Small increases in Toll signaling have massive costs to reproductive potential

Our results reaffirm the well-established trade-off between immunity and reproduction (Schwenke et al., 2016), but also emphasize the sensitivity of this relationship (Figure 5e). Specifically, injecting beetles with the smallest dose of *cactus* dsRNA led to just a four-fold increase in constitutive *cec-2* expression and a similarly modest increase in beetles surviving Bt infection but resulted in a severe reduction in reproductive output. This is not surprising, considering previously documented costs to Toll overexpression (Beramendi et al., 2005; Ryu et al., 2008; DiAngelo et al., 2009) and the role of Cactus as a direct inhibitor of Toll transcription factors, representing a potent and costly form of chronic immune activation. Is immune dysregulation always this costly? When the negative regulator *pirk-like* in *Aedes aegypti* is completely knocked-out, there is an approximately 30% increase in final survival probability to *E. coli* infection but only an approximately 52% decrease in female egg laying (M. Wang et al., 2022), representing a far less costly process to increase resistance. We are not aware of any additional studies that directly investigate these attributes of negative regulators on host reproduction, but we expect that such studies will reveal unique trade-off relationships (e.g., concave or exponential) compared to our results, which will provide key empirical data for theoreticians to incorporate real-life trade-offs and mechanistic detail into models of host-parasite and life history evolution.

Theoretical models of immune system evolution, for example, generally incorporate the cost of increased resistance to fitness traits like reproduction, growth, or mortality (Boots et al., 2009). To keep things simple, models tend to stay agnostic to the molecular underpinnings behind changes in immune output (Schmid-Hempel, 2003). They often assume small mutational steps that increase immune output and infection resistance, which coincides with equal reductions to fitness traits (Schmid-Hempel, 2003; Boots & Bowers, 2004; Best et al., 2014; Buckingham & Ashby, 2022). However, our results clearly show that the cost of increasing parasite resistance is not always linear. Rather, the trade-off between resistance and reproduction from elevated Toll signaling follows a convex decay form (Figure 5e). While this relationship certainly does not reflect outcomes across all immune genes, future models could benefit from assessing multiple trade-off shapes beyond linear forms (Hoyle et al., 2008; Jessup & Bohannan, 2008; Buckingham & Ashby, 2022). Furthermore, integration of additional empirical examples would enable models to better mirror real-life consequences of immune variation. Researchers could then manipulate environmental parameters as needed and superimpose the most influential trade-offs onto fitness landscapes (Jessup & Bohannan, 2008), potentially aiding in understanding phenomena such as the natural variation of macrophage activation, which modulates the trade-off between enhanced cancer survival and increased susceptibility to autoimmune diseases like rheumatoid arthritis (Buscher et al., 2017).

### Dose-dependent Cactus depletion regulates Toll signaling

We found that *cactus* depletion affects the expression of AMPs associated with both IMD (*def-2 & 3)* and Toll (*cec-2* and *def-3*) pathways. This is consistent with previous studies highlighting the cross-talk between the TFs Dif-Dorsal (Toll) and Relish (IMD) in *T. castaneum* (Altincicek et al., 2008; Yokoi, Koyama, Ito, et al., 2012; Yokoi, Koyama, Minakuchi, et al., 2012; Behrens et al., 2014; Tate & Graham, 2017; Yokoi et al., 2022). Pathway cross-talk is an increasingly appreciated feature of immune signaling across Insecta, including in the Hemipteran *Plautia stali* (Nishide et al., 2019) and Hymenopteran *Apis mellifera* (Lourenço et al., 2013) and has even been described in *D. melanogaster* (Tanji et al., 2007), indicating less pathway specificity than predicted by early studies in fruit flies (Lemaitre et al., 1997). At the same time, our data indicate the existence of distinct negative regulatory elements that differentially affect decay rates within these two pathways. Following *cactus* depletion, for example, the expression of *def-2* shows a delay in decay rate after microbial challenge but still ultimately resolves by 48 hours post exposure. Regardless of the magnitude of Toll activation, on the other hand, *cec-2* maintains maximum expression, suggesting the existence of IMD-specific negative regulators compensating for NF-κB dysregulation that do not compensate for Toll pathway-mediated effectors. In *D. melanogaster*, microbial challenges are also necessary for extracellular amidases, zinc finger proteins, and the JAK/STAT, JNK, and MAPK pathways to suppress the IMD pathway (F. Wang & Xia, 2018). It is intriguing to consider whether the IMD-specific decay we observe is driven by conserved mechanisms with *D. melanogaster* or novel regulatory factors. Such studies could shed light on conserved or novel mechanisms of immune modulation and provide a deeper understanding of genetic network robustness and resiliency, providing potential therapeutic targets for diseases characterized by overactive immune responses.

### Toll signaling increases circulating hemocytes after microbe challenge

Prior research in *D. melanogaster* larvae indicates that overactivation of Toll via *cactus* RNAi triggers hemocyte proliferation, melanotic tumors, and lamellocyte differentiation (Lemaitre et al., 1995; Qiu et al., 1998; Banerjee et al., 2019). In our study, we explored whether adult *T. castaneum* beetles exhibit similar proliferation from increased Toll signaling. Unlike *D. melanogaster* larvae, we observed no increase in circulating hemocytes prior to challenge. Interestingly, heightened Toll signaling did increase circulating hemocytes after microbial challenge, but not after septic wounding. This suggests a Toll-dependent hemocyte proliferation mechanism outside of the beetle wounding response that is enhanced by unrestricted Dif-Dorsal translocation. Our results also differ from a study in adult *Anopheles gambiae*, which found Toll overactivation did not significantly alter circulating hemocyte numbers two days post microbe challenge (Barletta et al., 2022). These disparate results highlight an interesting possibility that different mechanisms may regulate hemocyte proliferation and differentiation in these three model organisms.

### Increasing Toll signaling amplifies damage to host health

Toll overactivation results in severe costs to host health through neuromuscular destabilization (Beramendi et al., 2005), fat body depletion (DiAngelo & Birnbaum, 2009), and gut instability (Ryu et al., 2008). Similarly, our study revealed that the dysregulation of Toll signaling in beetles resulted in shorter lifespans, greater body mass depletion, and increased probability of gut barrier instability. While these results align with our expectations, the differences observed between the doses introduce intriguing questions regarding the underlying mechanisms and factors influencing these outcomes.

In our study, we found that beetles treated with *cactus* RNAi became increasingly lethargic until their early death. We also found that by mitigating Toll dysregulation with lower doses, lethargy is delayed, and death is less prevalent. Toll dysregulation in *D. melanogaster* also leads to lethargy, which is associated with destabilized neuromuscular junctions (NMJs) (Beramendi et al., 2005). NMJ disruption is linked to the alternative splicing of *dorsal* and *cactus*, which creates two distinct protein isoform pairs that mediate development and immunity (Dorsal-Cactus A) or NMJs (Dorsal-Cactus B) (Zhou et al., 2015). Surprisingly, Dorsal B, Pelle, and Cactus B work together, not in opposition, for proper NMJ functioning (Heckscher et al., 2007). Since our *cactus* RNAi targets both isoforms, subsequent research should evaluate whether the concentration of this particular Cactus isoform dictates the observed delayed phenotype, thereby improving our understanding of how organisms employ alternative splicing to circumvent costs from varying expression of pleiotropic genes (Williams et al., 2023).

To dramatically increase AMP production, *D. melanogaster* modifies its metabolic pathways by reducing insulin hormone levels, shifting from anabolic lipid metabolism to phospholipid synthesis and endoplasmic reticulum (ER) expansion (Martínez et al., 2020). This metabolic shift enhances triglyceride synthesis and lipid droplet formation while decreasing insulin signaling and hormonal levels (Cheon et al., 2006; Harsh et al., 2019; Suzawa et al., 2019). As a result, pupal triglyceride stores halve when Toll signaling is genetically induced (Martínez et al., 2020). Our findings suggest that even small levels of Toll activation can cause metabolic stress, resulting in lower body mass and egg production in female beetles. Dissecting the relationship between immune activation, metabolism, and egg production could offer crucial insights into immune-metabolic interplay and its evolutionary implications (Gupta et al., 2022).

Intestinal barrier dysfunction in *D. melanogaster* predicts metabolic defects in insulin signaling, shifts in immune signaling, and fly death more accurately than chronological age (Rera et al., 2012). This gut barrier instability is linked to excessive AMP expression, changes to the composition of gut microbiota composition, and stem cell hyper-proliferation, resulting in shorter fly lifespans (Ryu et al., 2008; Guo et al., 2014). Our study aligns with these findings, as we noticed significant gut instability from increased Toll signaling. We also observe that milder *cactus* depletion results in a lower incidence of destabilized guts. Unexpectedly, as the lower dosed beetles approached their mortality window, the proportion of dysfunctional guts did not increase, suggesting that even though gut dysfunction could contribute to mortality, it is not the primary driver of it. RNAi in *T. castaneum* depletes mRNA across all tissues (Posnien et al., 2009). While this complicates distinguishing gut immunopathology from other physiological dysfunctions, this is a challenge common to the field that requires further innovation to disentangle. Considering our findings, it would be interesting to explore whether gut instability is associated with AMP concentration, or if tolerance mechanisms effectively mitigate damage from immune signaling dysregulation at these low doses.

## Conclusion

Using our *cactus* RNAi experimental framework, we conducted a novel quantitative assessment of the benefits of increased immunity and the significant fitness costs incurred by immune dysregulation in the absence of infection (Figure S3). Specifically, we investigated the effects of increased Toll pathway signaling in *T. castaneum*, which resulted in increased AMP transcription and the number of circulating hemocytes leading to functional increases in resistance. By serially increasing the strength of Toll signaling, we demonstrated that the magnitude of Toll activation correlates with the ability of *T. castaneum* to resist and survive parasitic infection. Our findings reveal the costs of increased parasite resistance, as beetle lifespan, reproduction, mass, and gut integrity reduced as the degree of immune activation increased. Our results align with previously identified phenotypes from Toll pathway overactivation (Roth et al., 1991; Nicolas et al., 1998; Qiu et al., 1998; Beramendi et al., 2005; Anjum et al., 2013; Liu et al., 2016; Bingsohn et al., 2017; Germani et al., 2018; Sneed et al., 2022), but reveal the steep sensitivity of their relationship with immune activation. Our results stress the need for future empirical studies to adopt this type of experimental framework to quantify the benefits and costs of immune regulation.

## Materials and Methods

### Beetle rearing and experimental groups

For each experiment we put approximately 200 age-matched parental beetles on fresh beetle media (whole-wheat flour from MP Biomedicals and 5% yeast) to lay eggs for 24 hours and then we removed the parents onto new media. Once the eggs developed into pupae, we separated them by sex and transferred them to petri dishes (100 mm) containing either 100 unmated males or females with *ad libitum* media. We derived all experimental beetles from the ‘Snavely’ beetle population, originally collected from a Pennsylvania grain elevator in July 2013 and subsequently maintained in the laboratory since (Tate & Graham, 2015). We kept all beetles in a walk-in incubator at 30°C and 70% humidity.

### DsRNA synthesis

We designed the *T. castaneum cactus* primer sequences for dsRNA synthesis from the iBeetle Database (Ulrich et al., 2015) (Table S6). We then verified the primer pair for the absence of secondary effects by comparing phenotypes with a non-overlapping dsRNA fragment (Ulrich et al., 2015) and by blasting the primer pair against the new *T. castaneum* genome using NCBI’s Primer-BLAST (Ye et al., 2012). Next, we generated T7 promoter sequence-tagged DNA from *T. castaneum* cDNA via PCR using the Platinum Green Hot Start kit (Invitrogen). We then purified the PCR product using the QIAquick PCR Purification kit (Qiagen). Using the Megascript T7 kit (Invitrogen), we synthesized dsRNA overnight (Posnien et al., 2009). As a control for the induction of beetle RNAi, we used E. coli DNA as a template to produce dsRNA against a maltose binding protein E (*malE*) sequence (Yokoi, Koyama, Ito, et al., 2012). Finally, we quantified dsRNA concentration using the Qubit™ microRNA Assay Kit (Invitrogen).

### RNAi treatments

Recently, (Bingsohn et al., 2017) demonstrated that by diluting the dose of *cactus* dsRNA to as little as 0.001 nanograms, the RNAi induced mortality in adult *T. castaneum* was reduced from 100% to 75%, illustrating a system capable of quantitatively modulating RNAi-mediated phenotypes based on knockdown efficiency. For each experiment, we randomly assigned beetles to their respective dsRNA treatment groups and injected them with 0.5 uL of injection mixture. For the *cactus* RNAi group, we dissolved dsRNA in sterile insect saline at concentrations of 500, 50, or 5 ng/uL, while for the MalE control group, all injections used the 500 ng/uL concentration. We did not test gene expression for naïve beetles because our prior study showed no significant difference in constitutive expression between MalE-RNAi and naïve beetles (Jent et al., 2019). For injections, we followed the methods detailed in (Posnien et al., 2009). We applied the formula (2^−(ΔCt(*cactus*) – ΔCt(*malE*)^) * 100 across all time points to quantify RNAi treatment knockdown efficiency. By averaging these values, we determined the overall relative knockdown percentage for the entire experiment, with MalE ΔCt denoting the mean ΔCt for all MalE samples (Livak & Schmittgen, 2001).

## Microbial infections

### *Bacillus Thuringiensis* (Bt)

Bt, a sporulating, gram-positive, and obligate-killing bacterium, can infect insects through oral ingestion or septic injection, and once it gains access to insect hemolymph, it grows rapidly, causing death within 12-24 hours (Raymond et al., 2010; Nielsen-LeRoux et al., 2012). For all infections, we used the Berliner strain of Bt (ATCC 55177)(Jent et al., 2019). Two days after RNAi injections, we cultured Bt overnight for 12-15 hours from a glycerol stock at -80°C in Luria Broth at 30°C. We transferred 200 uL of the overnight culture to 3 mL of new LB for 1.5 hours. We diluted the overnight and log cultures to OD_600_ values of 1.0 and 0.5, respectively. We then combined 500 uL of each culture and centrifuged the mixture at 4°C and 5,000 rpm for five minutes. We removed the supernatant and washed the Bt pellet twice with one mL of sterile insect saline. We then re-suspended the washed Bt pellet in 150 uL of insect saline and diluted 1:20 with insect saline to obtain an LD 50 dose (5 x 10^8^ colony forming units (CFU)/mL). For the gene expression experiments, we inactivated Bt by heating to 90°C for 20 minutes. Beetles were infected by inserting an ultrafine insect pin dipped in the Bt mixture between the head and pronotum and were kept in individual wells in a 96-well plate after infection.

### Candida albicans

*Candida albicans* is a common opportunistic fungal parasite that has MAMPs typical of insect fungal parasites sensed by the Toll pathway. For our fungal challenge experiment, we used the *C. albicans* strain (ATCC 18804). Two days after RNAi injections, we cultured *C. albicans* overnight for 12-15 hours from a glycerol stock at -80°C in Luria Broth at 30°C. We diluted the overnight culture to an OD_600_ of 1.0. We then centrifuged one mL of the culture at 4°C and 5,000 rpm for five minutes. After washing twice, we re-suspended the fungal pellet in 150 uL of insect saline (1.89 x 10^8^ colony forming units (CFU)/mL) and heat-killed at 90°C for 20 minutes.

### Immune gene expression and Bt load quantification via RT-qpcr

To isolate RNA, we used the Qiagen RNeasy kit and eluted with 30 µL of nuclease-free water. For cDNA synthesis, we used 100-200 ng of RNA in a 5 µL reaction using the SuperScript IV VILO master mix (ThermoFisher Scientific). We then diluted the resulting cDNA with 40 µL of nuclease-free water. We conducted RT-qPCR using the PowerUp SYBR Green master mix (Applied Biosystems) on a Biosystems QuantStudio 6 Flex machine. The thermal cycling conditions consisted of an initial denaturation step at 95°C for 2 minutes, followed by 40 cycles of 95°C (15 seconds), 55°C (10 seconds), and 60°C (1 minute). We ran all samples in duplicate, and we used the average Ct value for subsequent analyses, if the technical replicates were within 1 Ct. If the technical replicates differed by more than 1 Ct, we repeated the reaction.

We selected immune genes for RT-qPCR analysis based on their previous correlation to Toll and IMD output in *T. castaneum*, as reported in a study by (Yokoi, Koyama, Minakuchi, et al., 2012). Specifically, we assayed the expression of *defensin-2* (TC010517), *defensin-3* (TC012469), and *cecropin-2* (TC030482) to represent the IMD, IMD and Toll combined, and Toll pathways, respectively (Herndon et al., 2020). To assess reference gene expression, we used the RPS18 primer pair provided in (Lord et al., 2010), which showed RPS18 expression is constant during fungal infection.

For all gene expression analyses, we used the Δct method to calculate relative expression values on a log2 scale for each gene in each sample by subtracting the mean ct value of the target gene from the reference mean ct value (Schmittgen & Livak, 2008). This method of data representation allows for easier comparisons of gene expression across timepoints than the ΔΔct method for fold change when the baseline treatment also changes (Critchlow et al., 2019). To analyze the effect of RNAi treatment on gene expression, we split the analysis into induction (0-6 hours) and decay (8-48 hours) stages, except for *cecropin-2*, where peak induction occurred at eight hours post-infection. We ensured normality of data by examining histograms and Q-Q plots of the standardized residuals. We performed linear modeling on gene expression using treatment, time, and their interaction as factors with the lme4 package in R (v. 4.3.0) and adjusted *p*-values for multiple comparisons using the Bonferroni method(Dunn, 1961). A significant effect for treatment indicates a difference in gene expression magnitude between treatments, while a significant effect for hour indicates differential gene induction or decay over time from microbial challenge. A significant interaction denotes a difference in induction or decay rates for the target gene in *cactus-*treated beetles relative to control-treated beetles. To investigate the impact of cactus RNAi dosage on the induction and decay of target AMPs, we employed a linear model (expression ∼ dose * hour), where “dose” is a continuous variable, and the MalE treatment group serves as a reference point with zero ng of *cactus* dsRNA. A significant effect for dose indicates a linear increase in the total transcription of the gene with increasing dose. A significant effect for hour denotes significant gene induction or decay. A significant interaction term indicates that the rate of induction or decay is dependent on *cactus* dsRNA dose. See Figure S4 for experimental design.

### Hemocyte proliferation quantification

We isolated total circulating blood cells (hemocytes) by perfusion bleeding the hemolymph in adult beetles. For this, we used a modified protocol from (King & Hillyer, 2013). Briefly, we made an incision through the lateral edge of the abdominal tergite segments V and VII using a feather blade, while the beetle was held dorsally with abdomen pointing downwards. We then inserted a glass microinjection needle into the thorax region and 50 μL of insect saline buffer was injected, and the diluted hemolymph that exited the posterior abdomen was collected onto one of the three 1-cm diameter etched rings on Rite-On glass slides (Fisher Scientific, Epredia). We perfused at a rate of 15-20 seconds per RNAi treated beetle, with the first 50 μL collected in one etched ring, followed by second and third in the other wells. We allowed the hemocytes to adhere to the slide for 20 min at room temperature and were fixed and stained for 20 min using 4% paraformaldehyde in phosphate buffer saline (1xPBS) and Hoechst 33342 nuclear stain (Invitrogen, Carlsbad, CA, USA) in phosphate buffered saline (PBS). We then mounted the slides using coverslips with Aqua Poly/Mount (Polysciences; Warrington, PA, USA). We captured images with a Nikon Ti-E inverted microscope and Zyla sCMOS digital camera, then we counted hemocytes using the NIS-Elements software (Nikon). We performed the experiment over a course of three biological blocks/trails, with each composed of 3-4 beetles per treatment combination. We aggregated data from all the trails. Given the non-normal distribution of residuals, we used a series of Wilcoxon rank sum tests for all combinations of treatment, time, and infection. We corrected for multiple tests using the False Discovery Rate (FDR) (Benjamini et al., 2001).

### Antibacterial activity

To measure AB activity, we closely followed the protocol from (Khan et al., 2016), modified from Roth et al., 2010 by measuring the inhibition zones produced by the whole-body homogenate of RNAi treated beetles on a lawn of Bt bacterial growth. To eliminate the confounding effects of quinones (secreted from the odoriferous defensive stink glands) on the AB activity, we froze beetles for 20 minutes at -80°C 3-days post RNAi treatment (N = 40-41 beetles per treatment & sex combinations). Freezing at -80°C triggers the release of quinones from the thorax and abdomen, which can otherwise prevent microbial growth and confound the results of the AB activity. We then cleaned the frozen beetles using 70% ethanol (surface sterilization) to remove any remaining quinones and washed twice with 100 μL insect saline. We prepared whole body homogenates of individual RNAi treated beetles using 30 μL sterile filtered Bis-Tris buffer (0.1 M, pH adjusted to 7.5, Sigma-Aldrich ltd) supplemented with 0.01% Phenylthiourea acid (PTU; Sigma-Aldrich ltd) to inhibit melanization. We then centrifuged the homogenate at 7500 rpm at 4°C for 10 minutes and separated the supernatant (on ice).

We prepared LB agar plates, using the overnight *Bt* culture in LB Broth at 30°C to an optical density of 1.0. Meanwhile, we separately autoclaved a LB broth with 1% agar and allowed it to cool in a water bath held at 45°C. Following this, we added the Bt bacterial culture to the agar medium and mixed by shaking, giving a final OD concentration of 0.001. We poured 6 mL of the mixture into each 75 mm Petri dish, with constant swirling to obtain homogenous distribution of bacterial culture. We then punched 4 wells (2x2 mm diameter) in each plate and added 4 µl homogenate from RNAi treated beetles to each of the wells. We added 2 µl Kanamycin (2 mg/ml) to every plate as positive control. We incubated the plates at 30°C for 16 h, at which point we measured the clear zones of inhibition of Bt bacterial growth around wells. We used the mean value by calculating the average of the horizontal and vertical diameters around each well as a proxy for the AB activity. Since the data is not normally distributed, we used a non-parametric Wilcoxon rank sum test to analyze the AB activity.

### Host survival experiments

For survival to parasite infection, we gave beetles an LD50 dose (6.5 x 10^^8^ CFUs/mL) of live Bt and monitored survival every 30 minutes for 12.5 hours and again at 24 hours post infection (N = 60-65/dose), We censored individuals that died at or before four hours post infection, since mortality most likely resulted from stab trauma. For estimating lifespan, we randomly assigned beetles to the four RNAi treatment groups (N = 30-32 beetles/dose, N = 15-16 beetles/sex), put them into individual wells in a 96-well plate with *ad libitum* media, and monitored them daily for 17 days. To compare survival among treatments for both experiments, we used weighted Cox Proportional Hazard tests (R package coxphw) to obtain a hazard ratio (Dunkler et al., 2018). The use of weighted Cox regression allowed us to obtain an average hazard ratio, despite our data not meeting the assumptions of the Cox proportional hazards model.

### Resistance to live Bt infection

To quantify Bt load during infection, we used the previously validated and published primers by (Tate & Graham, 2015, 2017) (Table S6), which correlate with bacterial CFU counts on agar plates. We performed a Kruskal-Wallis rank sum test to determine whether there were significant differences in Bt load for the four *cactus* dsRNA doses, since the data is non-parametric.

### Female reproductive output

We randomly assigned 7-day old unmated female beetles to one of four RNAi treatments, injected dsRNA, and allowed for 24 hours of recovery in individual wells in a 96-well plate with media. We paired females with a single aged-matched unmated male in a 2.0 mL centrifuge tube with 0.5 mL volume of media for 24 hours. We then separated the females and put them in their own petri dish with flour. After 24 hours, we counted the eggs for each individual and put the females on new flour. We allowed the eggs to develop for 21-days and counted the total number of viable larva for each female individual. We employed a zero-inflated negative binomial (ZINB) generalized linear model (R package MASS) to analyze the effect of dose and day on fecundity (fecundity ∼ dose*day), since the data is non-parametric and the ZINB model showed the lowest AIC score among all the tested glm types. We interpreted a significant effect for dose as indicating that the dosage of *cactus* dsRNA affected the number of eggs laid, while a significant effect for day indicated whether each day was unique from the others in terms of egg-laying behavior. A significant interaction term was interpreted as indicating whether the effect of dose on egg-laying changed over time.

### Body mass and fat body dissections

We cleaned female beetles twice by vortexing in 5 mL sterile insect saline and dried via Kimwipe. We measured average body mass changes on days three, four and five post RNAi injection. We then stored beetles in individual wells in a 96-well plate with beetle media until weighed. To analyze the changes in beetle mass, we used a linear model (mass ∼ dose + day). A significant main effect of dose indicates that the dosage of cactus dsRNA linearly changes the weight of female beetles. A significant effect of day indicates whether there are significant differences in weight among individual days.

### Gut/intestinal integrity quantification (Smurf assay)

We assessed gut barrier integrity of RNAi treated beetles by placing the beetles on blue food (n = 10-12 beetles per condition) prepared using 2.5% (w/v) FD&C blue dye no. 1 (Spectrum Chemicals) and sterilized wheat flour, prepared using the previously described protocol by (Rera et al., 2012). We examined the distribution of blue food dye after 24 h feeding. We randomly assigned beetles (females and males separately) to one of four RNAi treatment groups (*malE*-250 ng*, catcus*-0 ng, *cactus*-25 ng, *cactus*-250 ng) and held them in individual wells in a 96-well plate. On days three, five, and eight following RNAi treatment, we removed a subset of beetles from flour and starved them for four-six hours. We then put the starved beetles back on flour mixed with blue food dye. After 24 hours of exposure to blue food, we dissected each beetle by removing both wings and looked for any leakiness in the gut-intestine barrier and scored the beetles as blue “smurf” phenotype. We used a binomial generalized linear model (glm) (smurfs ∼ dose * day) to analyze the effect of *cactus* dsRNA dosage on gut integrity. A significant effect of dose indicates that the dosage of *cactus* dsRNA determines the number of smurf beetles, while a significant effect for day is interpreted as changes in gut integrity over time. A significant interaction indicates that the number of smurfs over time is affected by the dose of *cactus* dsRNA.

## Acknowledgements

We would like to thank Louise Perrier for her help defining the numerical function that fits the shape of our trade-off. We thank Morgan Pfeffer, Alissa Williams, Allyson Ray, Reese Martin, and Jakob Heiser for comments and discussion. This work was supported by the National Institute of General Medical Sciences at the National Institutes of Health (grant number R35GM138007 to A.T.T.).

## Data Accessibility

All data is included in the supplemental files.

## Author Contributions

A.T.T conceptualized and obtained funding for this study. J.T.C, A.P, and A.T.T designed the experiments. J.T.C, A.P, and K.Y.Z conducted the experiments. J.T.C analyzed the data. J.T.C and A.T.T wrote the manuscript. All authors critically reviewed the drafts and approved the final version for publication.

## Supplementary Tables and Figures

Table S1. Effect of cactus RNAi (250ng) on total circulating hemocytes

Table S2. Effect of cactus RNAi dose on flour beetle lifespan

Table S3. Effect of cactus RNAi dose on female flour beetle reproductive output

Table S4. Effect of cactus RNAi dose on female flour beetle body mass

Table S5. Effect of cactus RNAi dose on flour beetle gut integrity Table S6. Primer sequences

Figure S1. Unrestrained Toll signaling increases resistance to Bt infection.

Figure S2. Unchecked Toll signaling disrupts intestinal integrity.

Figure S3. Summary of results when assessing the benefits and costs of elevated Toll pathway signaling.

Figure S4. Experimental design.

