## Supplementary figures for "Mapping the functional form of the trade-off between infection resistance and reproductive fitness under dysregulated immune signaling"

Authors: J.T.C, A.P., K.Y.Z, and A.T.T.

Figure S1. Unrestrained Toll signaling increases resistance to Bt infection. To measure shifts in host resistance to bacterial infection from *cactus* (250 ng) RNAi treatment, beetles were given an LD-50 dose of Bt and sacrificed six hours later. Relative bacterial density for each individual within each dsRNA treatment was quantified via RT-qPCR and calculated as the difference between Bt-specific and host reference gene expression (RP18s) on a log2 scale ($p < 0.05$).

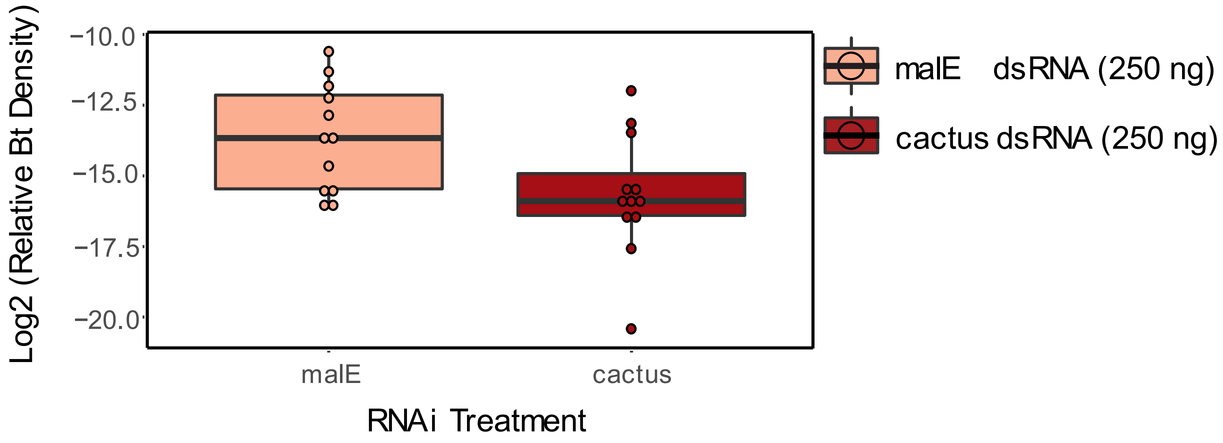

Figure S2. Unchecked Toll signaling disrupts intestinal integrity. Beetle gut integrity was measured by feeding adults flour stained blue and observing whether the blue dye entered the beetle hemolymph. Beetles with blue food dye in their hemolymph are counted as “smurfs”.

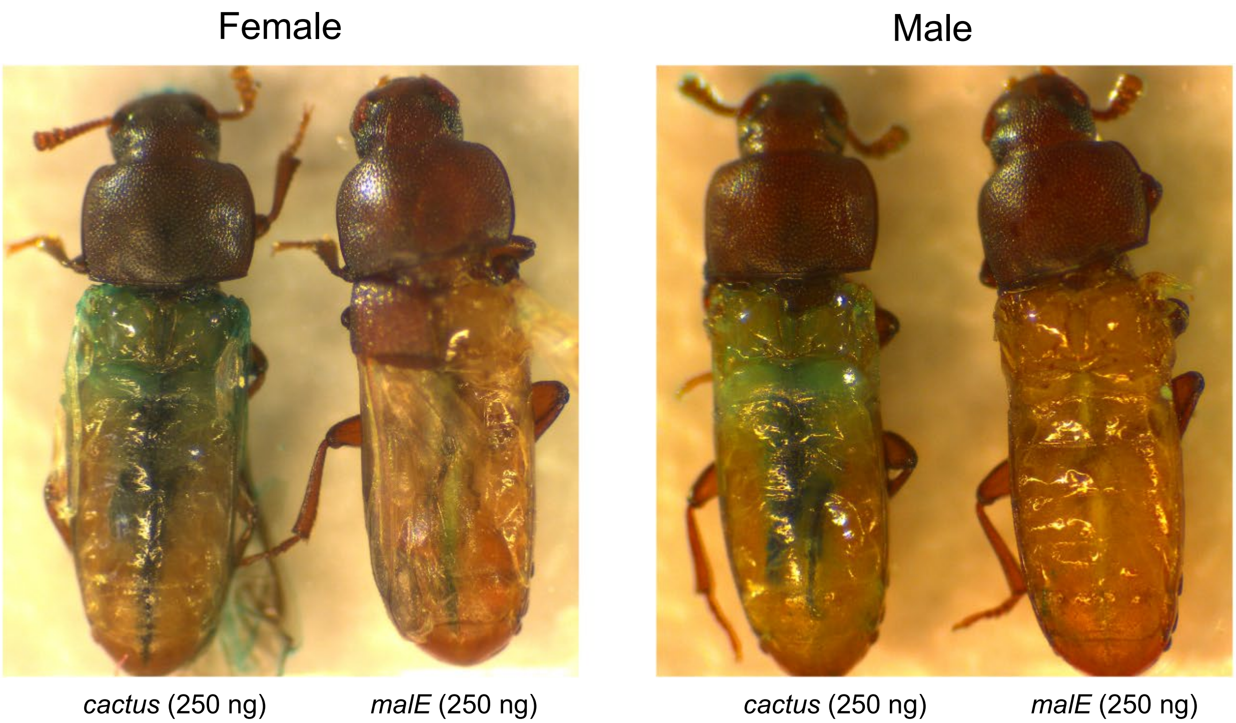

Figure S3. Summary of results when assessing the benefits and costs of elevated Toll pathway signaling.

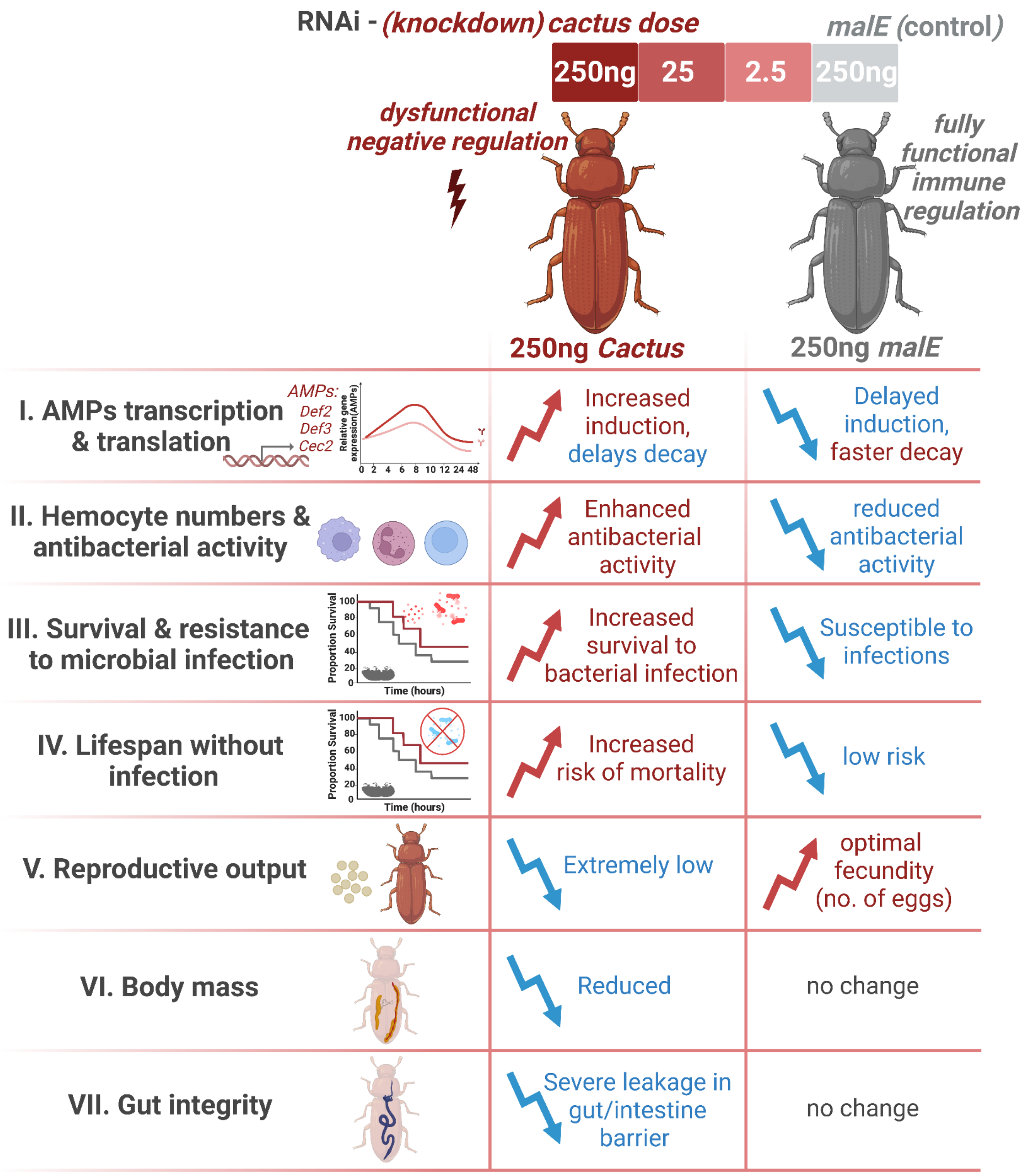

Figure S4. Experimental design. Our study investigated the benefits and costs of enhanced Toll signaling utilizing RNAi with four increasing concentrations of *cactus* dsRNA. **I)** Microbes *Bacillus thuringiensis* and *Candida albicans* were used to elicit immune activation. **II)** Immune gene expression and Bt load quantification were measured using RT-qPCR. **III)** Antibacterial activity was assessed by observing the inhibition of bacterial growth on a lawn of Bt. **IV)** Changes in total circulating hemocytes was examined by perfusing the hemolymph in adult beetles, staining, and counting the adhered hemocytes. **V)** Survival to pathogen infection was determined by monitoring RNAi treated beetles post-infection with an LD50 dose of Bt. **VI)** Female reproductive output was evaluated by pairing treated female beetles with male beetles for 24 hours and counting the number of eggs laid over three days. **VII)** Body mass and fat condition were assessed by cleaning and weighing beetles on the 3rd, 4th, and 5th day post-RNAi treatment. Additionally, on the 5th day, beetles were dissected and captured in images for fat body depletion. **VIII)** Gut integrity was examined by feeding the beetles media mixed with blue food dye and observing whether the dye entered the hemolymph.

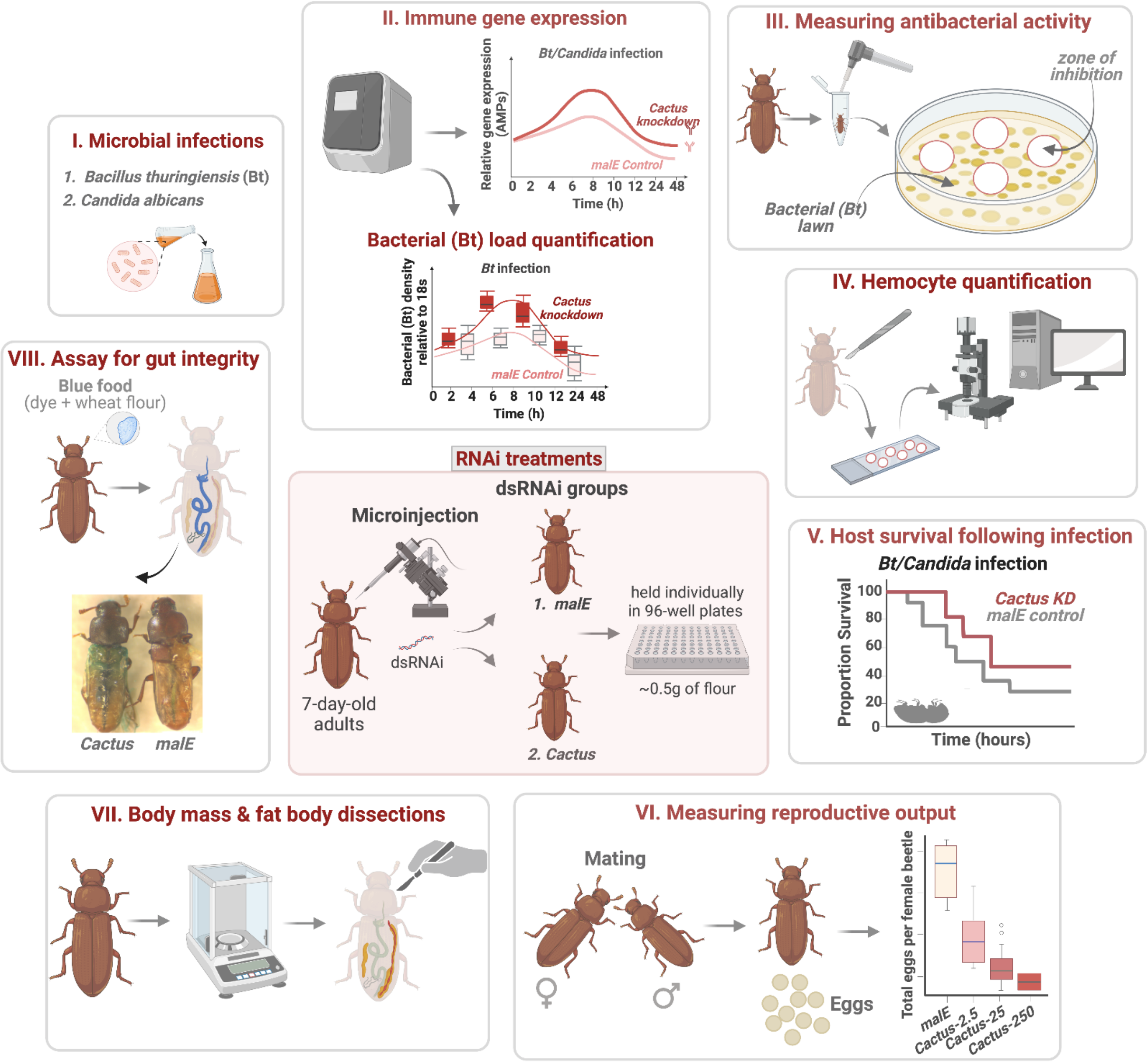
